# *Staphylococcus aureus* urease is controlled by a complex regulatory network and promotes dissemination during catheter-associated urinary tract infections (CAUTI)

**DOI:** 10.1101/2025.11.26.690714

**Authors:** Jana Gomez, Jesus Duran-Ramirez, Maristhela Alvarez, Shane Cristy, Jennifer N Walker

**Affiliations:** Department of Microbiology and Molecular Genetics, McGovern Medical School, Houston, TX, 77030, USA; The University of Texas MD Anderson Cancer Center UTHealth Houston Graduate School of Biomedical Sciences, Houston, TX, 77030, USA; Department of Epidemiology and Center for Infectious Diseases, UTHealth School of Public Health, University of Texas Health Science Center at Houston, Houston, TX, 77030, USA

## Abstract

Catheter-associated urinary tract infections (CAUTIs) are one of the most common hospital-associated infections in the United States, accounting for >1 million cases annually. One CAUTI-associated pathogen, *Staphylococcus aureus,* is commonly found persisting asymptomatically in the bladder of catheterized individuals, increasing these individuals’ risk of developing CAUTI. Importantly, *S. aureus* is not only associated with severe symptoms, including bacteremia and septic shock, but it also produces a common uropathogen associated virulence factor, urease. Despite its importance, urease has only been well-studied in the uropathogen *Proteus mirabilis.* While previous studies identified three *S. aureus* urease regulators, CodY, CcpA, and Agr, the environmental signals required for expression and activity, and the contribution of urease to CAUTI have not been explored. In this study, we investigate how the urease regulatory pathways coordinate expression and activity in response to environmental signals present within the catheterized urinary tract. Our findings demonstrate that stationary phase significantly induces *S. aureus* urease expression and activity. Additionally, we identified predicted binding sites in the urease promoter for three regulators previously implicated in urease expression – CodY, CcpA, and SigB – as well as SrrA – a previously unrecognized regulator of urease. A binding site for Agr was not present in the urease promoter. Expression and activity assays confirm the role of CodY, CcpA, SigB, SrrA, and Agr in regulating urease. Importantly, single nucleotide polymorphisms identified in the urease promoter of clinical isolates enhance urease expression. Additionally, urease promotes biofilm formation under catheterized urinary tract-like conditions in vitro and dissemination from the bladder to the kidneys at 1 day post infection in a mouse CAUTI model. Together, our data not only provide insight into the regulatory pathway controlling *S. aureus* urease but also emphasize the importance of studying these mechanisms in a model that mimics the host environment.

**AUTHOR SUMMARY:** Catheter-associated urinary tract infections (CAUTIs) are common hospital-associated infections and are caused by many different pathogens. Of these pathogens, *Staphylococcus aureus* is particularly problematic, as it often spreads to the bloodstream and results in septic shock. One of the primary virulence factors that is important for uropathogens is urease. Urease breaks down urea in urine, which results in crystalline structures that encrust urinary catheters and promote reflux to the kidney. Despite producing urease, little is known about how *S. aureus* regulates the enzyme. In this study, we investigate how regulatory pathways coordinate the expression and activity of urease in response to environmental signals. We show that growth during stationary phase and in conditions that mimic the urinary tract urease expression and activity are increased. We also show that five different regulators – CodY, CcpA, SigB, SrrA, and Agr control urease expression and activity. Additionally, genomic changes identified in the urease regulatory pathway of clinical urinary catheter-associated isolates enhance urease expression. Importantly, urease promotes biofilm formation and dissemination during CAUTI. Our study provides insight into the complex regulatory mechanisms controlling urease and highlights the role urease plays in the development and virulence of *S. aureus* CAUTI.

## INTRODUCTION

Of the more than 30 million urinary catheters placed each year, 10-30% become infected, which translates to over 1 million cases of catheter-associated urinary tract infection (CAUTI) annually^1–7^. The high infection rates combined with the frequency of catheter use has led to symptomatic CAUTIs becoming the most common hospital-associated infection in the United States. CAUTIs contribute to substantial morbidity and mortality and result up to $1.83 billion/year in associated preventable costs^8–11^. CAUTIs are also caused by a wide array of uropathogens, with no single pathogen accounting for more than 30% of infections^7–11^. Additionally, asymptomatic bacteriuria (ASB) – colonization of the bladder without symptoms – increases by 3-8% each day of catheterization, resulting in virtually all long-term urinary catheters (placed for >30 days) becoming colonized. Persistent ASB can increase the risk of developing a symptomatic CAUTI and act as a reservoir for multidrug resistant bacteria^8–12^. Furthermore, recent studies have indicated that there is a further shift in organisms causing ASB compared to CAUTI, with Gram-positive bacteria, including *Enterococcus faecalis* and *Staphylococcus aureus*, some of the most frequent colonizers^9,11^.

Of the uropathogens that commonly result in catheter-associated ASB, a majority produce the enzyme urease, including *P. mirabilis, Klebsiella pneumoniae,* and *Staphylococcus aureus*. Urease is a nickel-dependent metalloenzyme that hydrolyzes urea into ammonia and carbon dioxide, which result in an alkaline environment that promotes the formation of precipitates^13–16^. These precipitates can encrust urinary catheters and can contribute to catheter blockages, which lead to device failure and ultimately decrease the quality of life of individuals who are catheterized long-term as they require more frequent catheter changes^13,14,17^. Furthermore, catheter blockages can result in urine reflux to the kidneys, which increases the risk of secondary bacteremia^18,19^. Yet, the role of urease during the lifecycle of these uropathogens in the catheterized bladder has only been well studied in the context of *P. mirabilis* CAUTI and ASB, leaving the other urease-producers largely uncharacterized. Notably*, S. aureus* is second only to *P. mirabilis* in causing catheter blockages^13,20^, which suggests that *S. aureus* contributes substantially to device blockages and ultimately, decreased quality of life. Impactfully, *S. aureus* is often associated with more severe disease, including high rates of dissemination to the bloodstream from the catheterized bladder, which subsequently leads to increased rates of sepsis and shock compared to other uropathogens^11,21–23^.

Despite the severity of *S. aureus* CAUTI, few studies have investigated the pathogen’s lifecycle in the catheterized bladder or the contribution of urease to more severe patient outcomes. Previous studies investigating the regulatory pathways dictating *S. aureus* urease production demonstrated that several well characterized regulators control expression of the enzyme. These regulators include the accessory gene regulator (Agr) system – the quorum sensing system in *S. aureus*^24,25^, the transcriptional regulator CodY – a global regulator that responds to levels of branched chain amino acids (BCAA) and GTP in the environment^26^, and the carbon catabolite protein A (CcpA) – a carbon catabolite regulator that is activated in the presence of high levels of glucose in the environment^27,28^. Specifically, while mutants of *ccpA* and *agr* result in lower urease expression, a mutation in *codY* results in higher urease expression compared to the parent strains – demonstrating that these regulators activate and inhibit urease, respectively^26,28^. Additionally, the sigma factor B (SigB) global regulator has been implicated in urease regulation by the identification of a predicted transcription start site upstream of the urease operon ^29^. Yet, SigB has not been genetically confirmed to regulate urease^29^. Despite these studies, there has been no thorough investigation of the regulators involved in *S. aureus* urease expression and activity or the environmental signals that may be important for this regulation in the catheterized bladder.

In this study, we characterized the regulatory pathways controlling urease and determined the enzyme’s role in *S. aureus* biofilm and CAUTI. Our data indicate that growth stage and environmental conditions influence urease expression. Specifically, stationary phase growth in urine-like conditions induces the highest levels of urease expression and activity. Additionally, consensus binding sequences of CodY, CcpA, SigB, and two additional response regulators, SaeR and SrrA, were identified in the urease promoter, suggesting these directly regulate urease. Characterization of these predicted regulators demonstrate that urease expression and activity are inhibited by CodY and SrrA and activated by CcpA, SigB, and Agr. A mutation in SaeR did not have a significant effect on urease expression or activity under the conditions tested.

Impactfully, bioinformatic analyses of the urease promoter among urinary catheter-associated clinical isolates identified a common single nucleotide polymorphism (SNP) within the predicted consensus binding site of SigB that differed from the well characterized strain JE2 (parent strain isolated from an skin abscess)^12,30,31^. Additional studies demonstrated that this SNP resulted in a significant increase in urease expression in a urinary catheter-associated isolate compared to the JE2 promoter in the same background. Furthermore, in contrast to other studies performed in rich media, our results show that under conditions that mimic the catheterized urinary tract, *S. aureus* urease promotes biofilm formation *in vitro*. Finally, results from a CAUTI mouse model demonstrate that urease is essential for dissemination from the bladder to the upper urinary tract, which supports clinical observations of high rates of secondary bacteremia associated with *S. aureus* CAUTI^21–23^. These findings not only untangle the complex regulatory network dictating urease expression, which is required to most efficiently coordinate its expression in response to specific signals, but also show the importance of urease in *S. aureus* biofilm formation and infection in the catheterized urinary tract.

## RESULTS

### Growth phase is important for urease expression

To determine whether *S. aureus* senses signals within the urinary tract environment to coordinate the expression of urease, a transcriptional reporter plasmid with the 200- base pair region upstream of the urease operon (i.e. the urease promoter) fused to GFP (pUre1) was generated in the JE2 background strain (**Figure S1A, Supplemental File 2**). The pUre1 plasmid did not alter the growth of these cells compared to the parent strain in rich or minimal media (**Figure S1B, S1C)**. Urease expression was then assessed across a 12-hour (hr) time course in various media, including a nutrient rich growth media (BHI) and low nutrient, urinary tract-like media conditions, such as artificial urine media (AUM), female human urine (HU), and urease broth (UB) (**Figure 1A, S2A, S2B)**. Strikingly, the stationary phase of growth displayed the most consistent impact on urease expression, where regardless of media, there was significantly higher urease expression compared to the logarithmic phase of growth (**Figure 1A, S2A, S2B)**. Furthermore, urease expression was significantly higher in HU compared to BHI and AUM. However, HU displayed high variation across biological and technical replicates. Together, these data suggest urease is most highly expressed during stationary phase under conditions that are present in the urinary tract.

**Figure 1:**
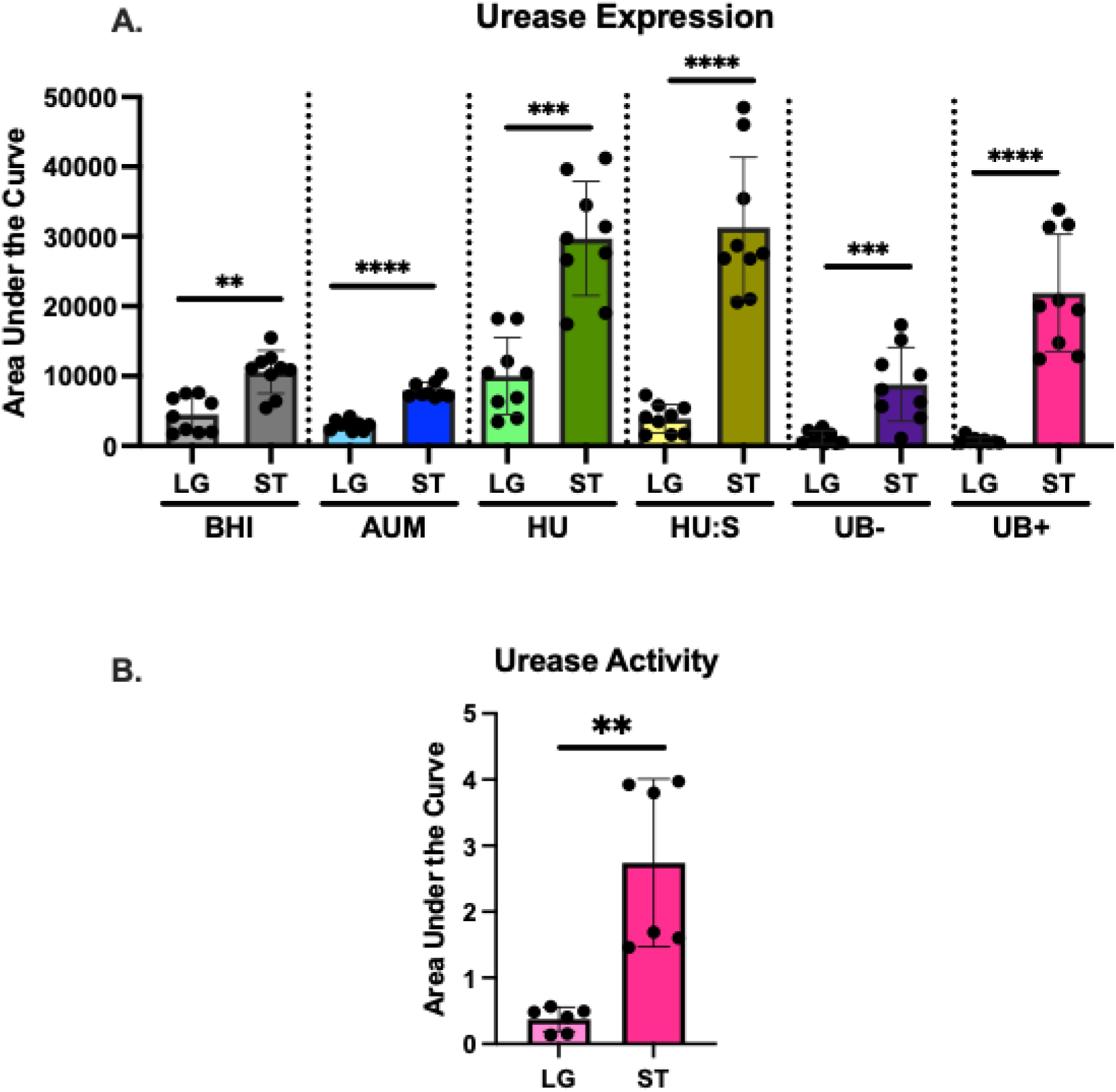
Urease expression and activity. A) Urease was analyzed for expression in BHI, AUM, female human urine (HU), a 1:1 ratio of HU and saline (HU:S), urease broth without urea (UB-), and urease broth with urea (UB+). Expression was analyzed in logarithmic (LG) and stationary (ST) phase cultures. Fluorescence was normalized to the fluorescence reading of the background strain that did not contain the GFP reporter plasmid. Each dot represents a technical replicate, with 3 technical and 3 biological replicates performed. Bars represent the mean of all replicates with the standard deviation. The area under the curve for each strain is quantified and significance was determined using the Mann-Whitney U test. B) Urease activity was assessed in UB. Urease expression and activity were measured over a 12-hour time course32. Each dot represents a technical replicate. A total of 2 biological with 3 technical replicates were performed. Bars represent the mean of all replicates with the standard deviation. The area under the curve for each strain is quantified and significance was determined using the Mann-Whitney U test.

### Growth phase is important for urease activity

In addition to expression, we also assessed whether growth phase impacts urease activity^32^. Activity was assessed across a 12-hour (hr) time course in urease broth (UB), as previously described^32^. UB contains a pH indicator that indirectly measures activity by detecting the increase in environmental pH that results from the breakdown of urea in the media by urease^32^. Consistent with urease expression, activity was also significantly higher in stationary phase compared to logarithmic phase (**Figure 1B, S2C**), emphasizing the importance of stationary phase growth in regulating activity.

Together, these data indicate that as *S. aureus* enters stationary phase, both urease activity and expression are upregulated.

### Identification of *S. aureus* urease regulators

To identify potential regulators responsible for sensing and coordinating the expression of urease, the urease promoter region was analyzed for the presence of previously published regulator consensus binding sequences^33–38^. Consistent with a previous report, two potential transcription start sites for the urease operon, one under the control of the primary sigma factor SigA and another under the control of alternative sigma factor SigB^29^, were identified (**Figure 2A, 2B**). Additionally, five consensus binding sequences were identified, including those for the previously identified urease regulators CodY^39^ and CcpA^27,39^, as well as one for SigB and two additional two component system response regulators SaeR and SrrA. Despite previous work indicating that the Agr quorum sensing system is involved in urease expression^39^, we did not identify a consensus binding site in the urease promoter for AgrA (**Figure 2A, 2B**). These results suggest that Agr regulates urease through an indirect mechanism, which is not uncommon for the quorum sensing system^24,40^. Overall, the identification of the five consensus binding sequences in the urease promoter suggests that CodY, CcpA, SigB, SaeR, and SrrA all regulate urease expression and activity.

**Figure 2:**
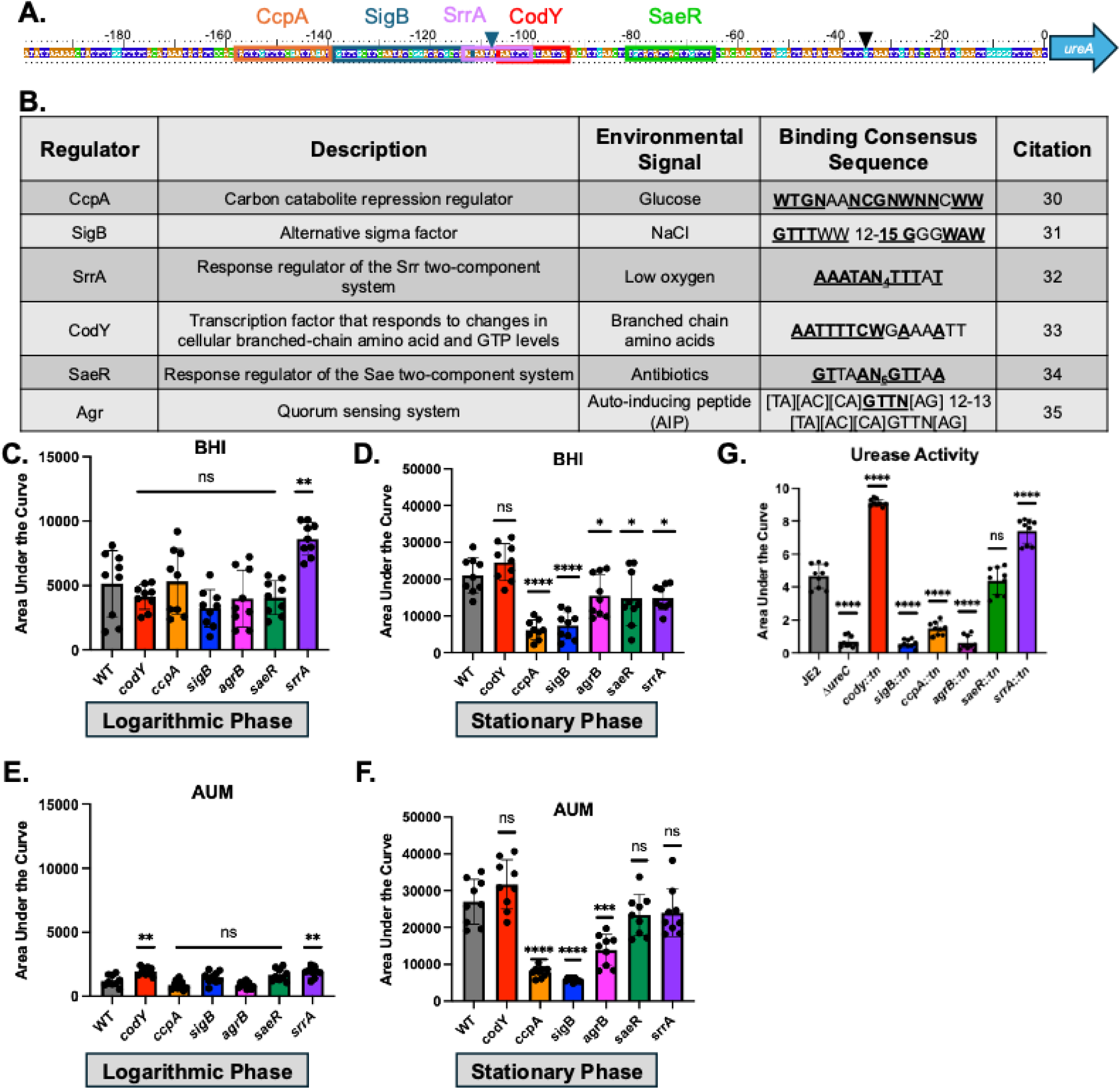
CcpA, SigB, SrrA, CodY, and Agr regulate S. aureus urease expression and activity. A) 200bp region upstream of the urease operon in JE2. Predicted binding sites for transcriptional regulators are denoted in boxes. Transcriptional start sites for the predicted SigA-controlled transcript (black) and the predicted SigB-controlled transcript (blue) are denoted by triangles29. B) Table detailing features of each predicted regulator of urease. Binding consensus sequences and their references are listed. Bolded and underlined characters indicate an exact match within the JE2 urease promoter. W indicates either an A or a T. N indicates any nucleotide. Urease expression in regulator mutants in the JE2 background at both logarithmic (C and E) and stationary (D and F) phase, in either BHI (C and D) or AUM (E and F) over a 12 hr time course. Fluorescence was normalized to the fluorescence reading of the background strain that did not contain the GFP reporter plasmid. Each dot represents a technical replicate, with 3 technical and 3 biological replicates performed. Bars represent the mean of all replicates with the standard deviation. The area under the curve for each strain was quantified and significance was determined using the Mann-Whitney U test. G) Urease activity in regulator mutants in the JE2 background. Urease activity was measured over a 12-hour time course32. Each dot represents a technical replicate, with 3 technical and 3 biological replicates performed. Bars represent the mean of all replicates with the standard deviation. The area under the curve for each strain is quantified and significance was determined using the Mann-Whitney U test.

### Agr, CcpA, CodY, SigB, and SrrA control urease expression

To determine the role of these predicted regulators in urease expression, loss of function mutations in each regulator were generated in the JE2 background strain (**Figure S3A**). AUM was prioritized for further urease expression experiments as it displayed a similar, yet more consistent, expression phenotype across biological and technical replicates compared to HU. To confirm mutations in each regulator did not affect growth, strains were assessed for growth in BHI and AUM (**Figure S2B-E**).

Compared to JE2, the *ureC* mutation grew similarly in BHI and AUM. However, the regulator transposon mutants displayed small but significant differences in growth compared to JE2. In BHI, the *codY, sigB, saeR,* and *srrA* mutants exhibited increased growth compared to JE2, while the *ccpA* and *agrB* mutants displayed a small growth defect (**Figure S3B, S3C**). In AUM, the *codY* and *sigB* mutants exhibited increased growth compared to JE2, while the *ccpA, agrB, saeR,* and *srrA* mutants displayed a small growth defect (**Figure S3D, S3E**). To determine the impact of each regulator on urease expression, the pUre1 plasmid was moved into each mutant background strain. Next, urease expression was measured in both BHI and AUM (**Figure 2C-F, S4A-D**). In BHI, when urease expression is low (e.g. logarithmic phase), compared to JE2 all the mutants displayed similar expression, except *srrA::tn,* which exhibited significantly higher expression (**Figure 2C, S4A**). However, in BHI, when urease expression is high (e.g. stationary phase), compared to JE2 all the mutants displayed significantly lower urease expression, except *codY::tn*, which exhibited similar expression to the wild type (**Figure 2D, S4B**). In contrast, in AUM, when urease expression is low (e.g. logarithmic phase), compared to JE2 both *codY::tn* and *srrA:tn* displayed significantly higher urease expression, while *ccpA::tn, sigB::tn, agrB::tn,* and *saeR::tn* displayed similar patterns of expression (**Figure 2E, S4C**). In AUM, when urease expression is high (e.g. stationary phase), *ccpA::tn*, *sigB::tn*, and *agrB::tn* exhibited a significant decrease in expression, while *codY::tn, srrA::tn,* and *saeR::tn* displayed similar expression (**Figure 2F, S4D**).

Additionally, to account for any differences in growth of the transposon mutant strains compared to JE2, fluorescence per cell was also calculated (**Figure S4E-H**). This analysis showed similar patterns for each strain compared to the analysis of each reporter strain normalized to its respective background strain (**Figure 2**). Overall, these results indicate that CcpA, SigB, and AgrB are activators, while CodY and SrrA act as inhibitors of urease expression.

### Agr, CcpA, CodY, SigB, and SrrA control urease activity

To determine the role of the six regulators on urease activity, the loss of function mutants in each regulator were assessed in the urease activity assay. Since our previously published urease activity data indicates that there is no detectable urease activity in the logarithmic phase of growth ^32^, activity assays were performed with stationary phase cultures. Compared to JE2, significantly higher urease activity was observed in *codY::tn* and *srrA::tn* mutant strains, while significantly lower urease activity was observed in the *sigB, ccpA,* and *agrB* mutant strains (**Figure 2G**). The *saeR* mutant did not have any effect on urease activity under these conditions (**Figure 2G, S4I**).

These data demonstrate that CodY and SrrA are repressors of urease activity, while SigB, CcpA, and AgrB are activators of urease activity. Importantly, these urease activity data are consistent with expression results^26,28^, further supporting the role of CodY, CcpA, SigB, Agr, and SrrA in regulating urease expression and activity.

### Identification of environmental signals that induce urease expression

The identified urease regulators are known to respond to specific environmental signals, including glucose (CcpA)^41^, NaCl (SigB)^42^, BCAA (CodY)^43^, low oxygen (SrrA)^44^, antibiotics (SaeR)^45^, and quorum sensing peptides (Agr)^46^. To determine whether these environmental signals impact urease expression, strains were assessed in AUM supplemented with each signal known to activate the respective regulator (**Figure 3**). During logarithmic phase, when *S. aureus* is actively utilizing glucose for growth, urease expression significantly increased in AUM supplemented with glucose in a dose dependent response compared to AUM alone (**Figure 3A, S5A**). However, urease expression was unaffected by the addition of glucose during stationary phase, as expected, as the bacteria are no longer utilizing glucose to grow (**Figure 3B, S5B**). Furthermore, this phenotype was partially ablated in the *ccpA::tn* mutant, indicating that this increase in expression is mediated by CcpA. Likewise, the addition of 0.5M NaCl significantly increased urease expression during logarithmic phase compared to AUM alone (**Figure 3C, S5C**), while there was no effect during stationary phase (**Figure 3D, S5D**). This increase in expression in logarithmic phase was abolished in the *sigB::tn* mutant background, indicating that this phenotype was SigB*-*mediated.

**Figure 3:**
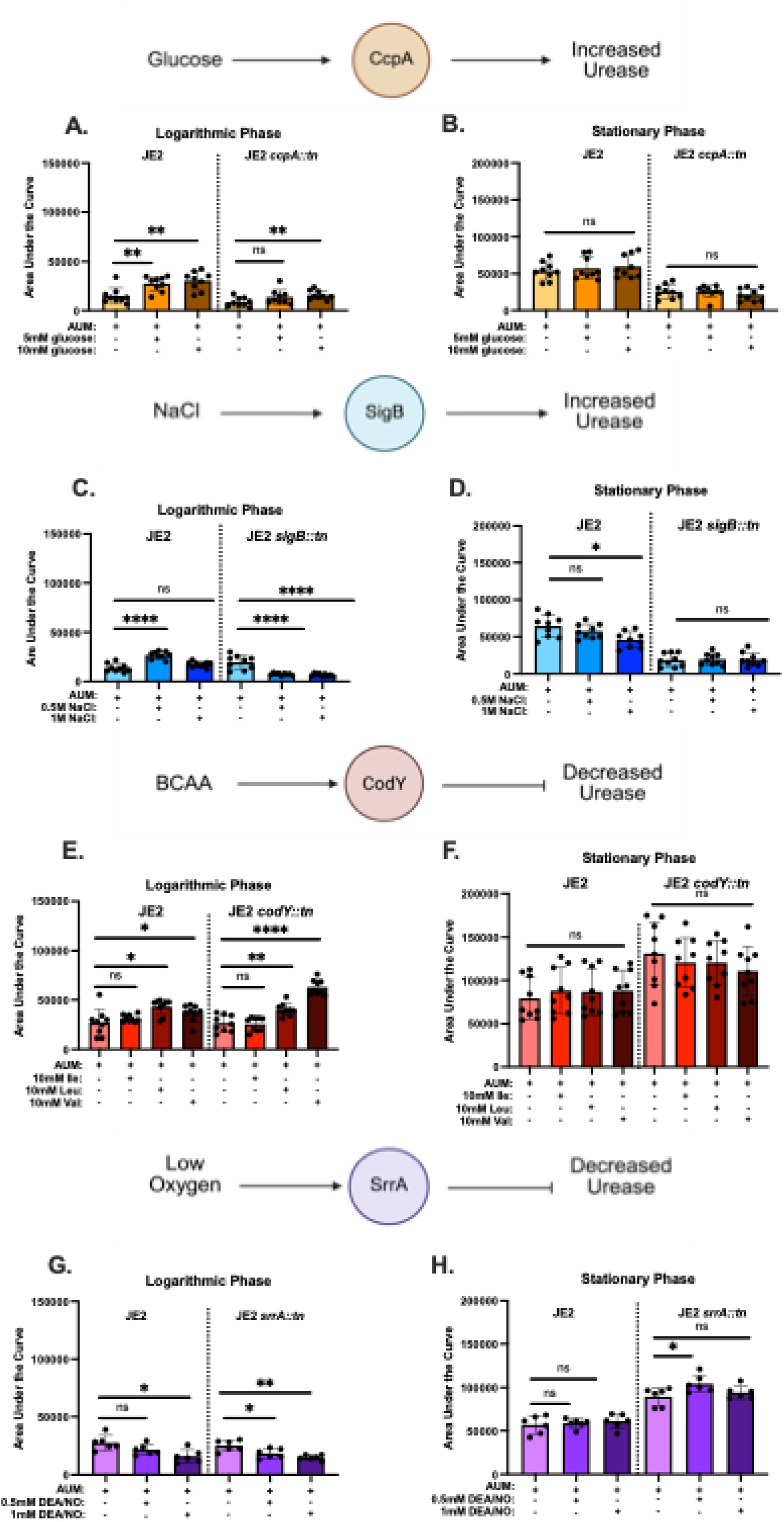
The impact of environmental signals on urease expression. Cells were grown in the presence of the CcpA signal glucose (A,B), the SigB signal NaCl (C,D), the CodY signal branched chain amino acids (E,F), or the SrrA signal DEA/NO (G,H) in cells in the logarithmic (A,C,E,G) or stationary (B,D,F,H) phase. Schematics indicating how each signal are expected to influence urease expression are depicted above each graph. Fluorescence was normalized to the fluorescence reading of the background strain that did not contain the GFP reporter plasmid. Each dot represents a technical replicate, with at least 3 technical and 2 biological replicates performed. Bars represent the mean of all replicates with the standard deviation. The area under the curve for each strain was quantified and compared for significance using the Mann-Whitney U test.

The addition of BCAAs (10mM of leucine and 10mM of valine) during logarithmic phase significantly increased urease expression compared to AUM alone (**Figure 3E, S5E**), while no change in urease expression was observed in stationary phase (**Figure 3F, S5F**). However, this phenotype was seen in both the JE2 and the *codY::tn* mutant background strains, indicating the observed increases in expression are likely independent of CodY. Similarly, during logarithmic phase, the addition of a nitrous oxide donator (DEA/NO) to AUM significantly reduced urease expression (**Figure 3G, S5G**), while there was there was no significant difference in urease expression in stationary phase (**Figure 3H, S5H**). However, both the JE2 and *srrA::tn* strains displayed a similar phenotype in both conditions, suggesting an SrrA-independent mechanism of regulation for DEA/NO. Additionally, during logarithmic phase, AUM was supplemented with supernatant from JE2 (positive control) or the *agrB::tn* (negative control)^46^. JE2 supernatant significantly increased urease expression in a dose dependent manner in JE2 in both the logarithmic and stationary phase (**Figure S6A-D**). This phenotype was also observed with the addition of JE2 supernatant to the JE2 *agrB::tn* strain during logarithmic and stationary phase. Addition of *agrB:tn* supernatant to *agrB:tn* in logarithmic phase and JE2 in stationary phase also induced urease expression, suggesting that the addition of supernatant, independent of the presence of auto- inducing peptide, is activating urease expression. Finally, during logarithmic phase, the addition of vancomycin significantly decreased urease expression compared to AUM alone (**Figure S6E, G)**. However, in stationary phase, the addition of vancomycin did not significantly alter urease expression (**Figure S6F, H**). This decrease in expression in logarithmic phase was observed in both the JE2 and *saeR::tn* mutant backgrounds, suggesting this phenotype occurs through an SaeR-independent mechanism. Together, these results identify several environmental signals, including glucose, salt, BCAA, low oxygen, and antibiotics, that influence urease expression. Notably, the quantities of these signals fluctuate in the urine, suggesting that urease expression is highly dependent on the composition of the urinary tract.

### Urinary catheter-associated isolates contain SNPs in the urease promoter

Our previous study determined that the most common *S. aureus* lineage asymptomatically colonizing the urinary tract was clonal complex (CC) 5^12^, which contrasts with JE2 – an CC8^47^. Furthermore, our previous report demonstrated that longitudinal *S. aureus* urinary catheter-associated isolates displayed increased urease activity over time^32^. To begin to investigate the role of the regulatory pathway in dictating urease expression and activity in clinical isolates, we analyzed the sequence conservation of the urease promoter among 171 urinary catheter-associated clinical *S. aureus* isolates (**Supplemental File 2**), along with JE2. Of the 172 sequences analyzed, the urease promotor was highly conserved among all the isolates. Clustering at 100% nucleotide identity revealed 11 unique sequence groups (**Figure S7A**). Strikingly, most isolates belonged to just two groups, comprising 67 (39%) and 49 (28%) isolates, respectively. However, the JE2 promoter was not represented within either of these two groups and instead grouped with only a single clinical isolate **(Figure S7A**).

The most common promoter sequence of the urinary catheter-associated isolates (pUre2) differed by two SNPs compared the JE2 promoter sequence (pUre1). Notably, one SNP was located in the predicted SigB consensus binding site (**Figure 4A**). This finding suggests that SNPs in the urease promoter may affect urease expression and activity in the catheterized bladder.

**Figure 4:**
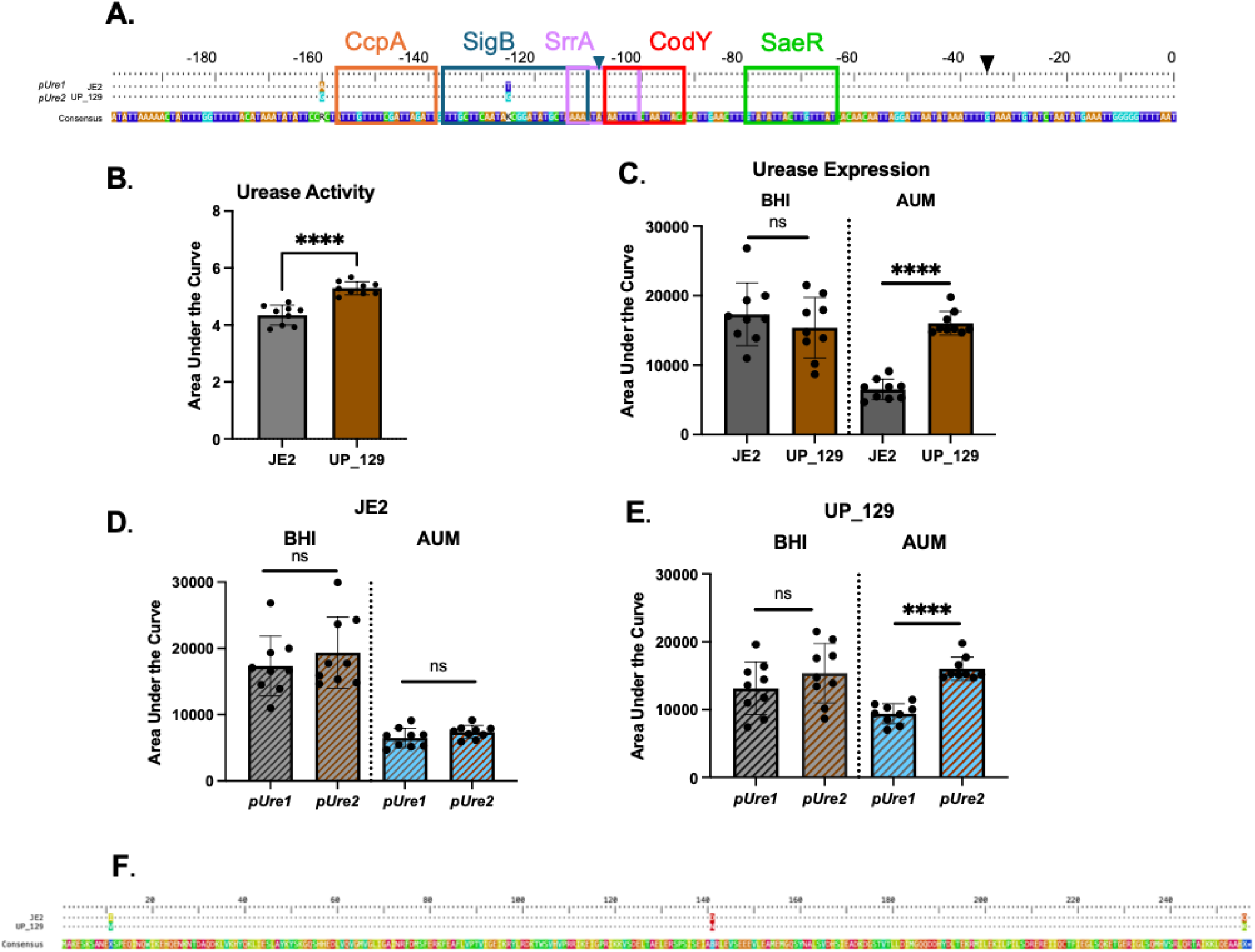
Clinically observed SNPs in the promoter alter urease. A) Comparison of the sequence of the upstream region of urease between JE2 and UP 129. Dots indicate a conserved nucleotide with the consensus. Predicted binding sites for transcriptional regulators CcpA, SigB, SrrA, CodY, and SaeR are denoted in boxes. Transcriptional start sites for the predicted SigA-controlled transcript (black) and the predicted SigB-controlled transcript (blue) are denoted by triangles29. Urease activity B) and urease expression (C) of JE2 compared to the clinical isolate UP 129. Each dot represents a single replicate for JE2 (black bars) or UP 129 (brown bars). Expression was analyzed in both BHI and AUM. Fluorescence was normalized to the background strain (JE2 or UP 129) without the fluorescent reporter. Promoter swap GFP reporter assays in the JE2 (D) and UP 129 (E) background. Fluorescence was normalized to the background strain (JE2 or UP 129) with no fluorescent reporter in BHI (gray) and AUM (blue). Gray bars indicate the native JE2 pUre sequence (pUre1) and brown bars indicate the native UP 129 pUre sequence (pUre2). Each dot represents a single replicate for each strain. A total of 3 technical and 3 biological replicates were performed for each experiment. Bars represent the mean of all replicates with the standard deviation. Area under the curve analysis was performed for each replicate and significance was determined using the Mann-Whitney U test. F) Alignment of the amino acid sequence of SigB in the JE2 (top) and UP 129 (bottom) strains. Dots indicate a conserved amino acid with the consensus. All alignments were created with the DECIPHER R package81.

### SNPs in the urease promoter of clinical isolates affect urease expression

To determine the importance of the SNPs within the most common urease promoter observed in the urinary catheter-associated isolates, the representative isolate UP 129 was selected for further characterization. UP 129 is not only a CC5 strain^30^, but it also contains the most common urease promoter sequence (**Figure 4A, S7A**). Impactfully, UP 129 also displayed significantly higher urease activity compared to JE2 (**Figure 4B, S7B)**. To determine if urease expression differed between the two strains, a promoter fusion using our GFP reporter assay was generated in UP 129 (**Figure S7C, Supplemental File 1)** and urease expression was assessed in BHI and AUM over a 12- hour time course. In BHI, there was no difference in the expression of urease between JE2 and UP 129 (**Figure 4C, S7D**). However, in AUM, UP 129 displayed significantly higher urease expression compared to JE2 **(Figure 4C, S7E**). Next, to determine the effect of the SNPs within the most common promoter on expression in each strain background, we generated promoter swaps using the GFP reporter assays generating pUre1 and pUre2 promoters in the opposite strain backgrounds. There was no significant difference between the expression of either JE2 pUre1 and JE2 pUre2 in BHI or AUM (**Figure 4D, S7F**). Similarly, there was no difference in the expression of UP 129 pUre1 and UP 129 pUre2 in BHI (**Figure 4E, S7G**). However, in AUM, UP 129 pUre2 resulted in significantly higher urease expression compared to UP 129 pUre1, suggesting that in this strain background the pUre2 promoter is more responsive to the urinary tract-like environment. The differences in expression of the pUre2 promoter between the JE2 and UP 129 backgrounds led us to compare the genomic conservation of the urease operon and the regulators between these two isolates. The protein sequences for *ureA, ureD, ureE,* and *ureG* were completely conserved between the JE2 and UP 129 strains, while the sequences of *ureB, ureC,* and *ureF* differed by only one amino acid between the two strains, indicating a high level of conservation of the urease operon (**Figure S8A**). Next, analyses of the animo acid sequences of the urease regulators, CodY, CcpA, AgrA, and SrrA, determined that the protein sequences are completely conserved between the JE2 and UP 129 strains (**Figure S8B**). However, the amino acid sequence of SigB differed in 3 different locations – amino acids 11, 141, and 246 (**Figure 4F**). These amino acid changes, in combination with the SNPs in the pUre2 sequence, may be additive and result in the observed increase in urease expression and activity in the UP 129 strain in AUM. Impactfully, these data demonstrate that as few as two SNPs within the promoter region of urease can have significant impacts on expression.

### Urease contributes to biofilm formation under conditions that mimic the catheterized urinary tract

Previous studies indicate that the genes in the urease operon are upregulated in biofilms^48^. As such, to determine the role of urease in biofilm formation under in vitro conditions that mimic the urinary tract environment, isogenic *ureC* mutants were generated in the JE2 and UP 129 backgrounds (**Figure S9A**). Biomass was measured after overnight incubation in BHI and AUM. Consistent with previous reports^49^, there was no significant difference in biofilm formation in either BHI or AUM between either of the parent or *ureC* mutant strains (**Figure 5A, 5B**). However, since it is well established that the healthy urinary tract differs from the catheterized urinary tract environment, as additional host proteins, including fibrinogen and serum albumin, are recruited and accumulate during catheterization, biofilms were also formed in the presence of human plasma (HP)^50,51^, which mimics the presence of these host factors^52^. As expected, the presence of HP significantly enhanced biofilm formation in all conditions (p-value less than or equal to 0.0188). Importantly, both JE2 *ΔureC* and UP 129 *ΔureC* formed significantly less biomass compared to their respective parent strains in AUM, but not in BHI (**Figure 5C, 5D**). These results indicate that urease contributes to *S. aureus* biofilm formation under conditions that mimic the catheterized urinary tract. Furthermore, these results emphasize the necessity of studying *S. aureus* urease under conditions that accurately represent the host environment.

**Figure 5:**
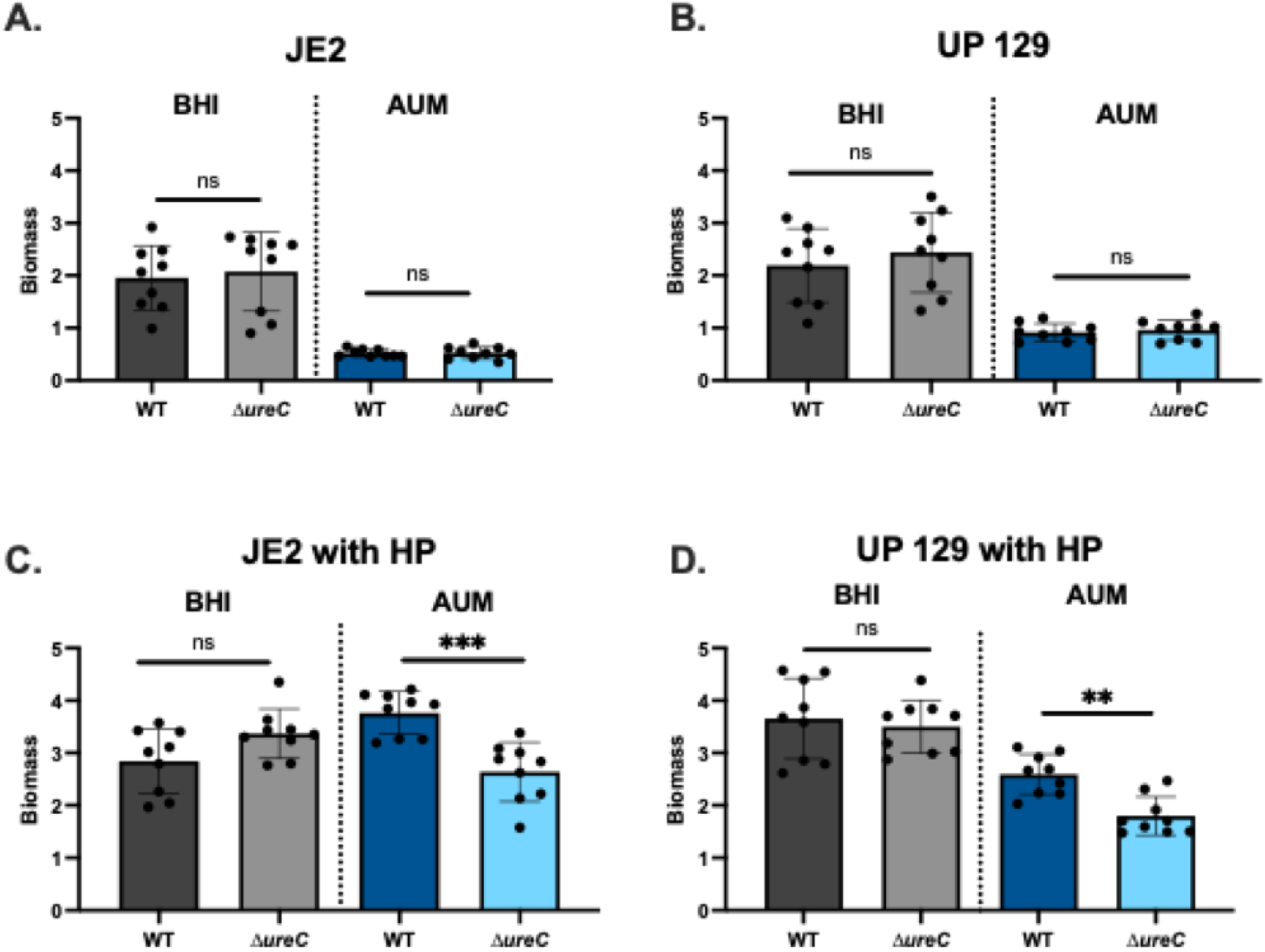
Crystal violet biofilms of WT and ΔureC strains. The biofilm biomass was measured under in vitro conditions in the JE2 wild type strain and the JE2 ΔureC strain (A) and the UP 129 wild type strain and the UP 129 ΔureC strain (B). To mimic the catheterized bladder environment, the biofilm assays were repeated in the presence of human plasma (HP) in both the JE2 (C) and the UP 129 (D) background. All assays were performed in BHI (gray bars) and AUM (blue bars). Each dot represents a technical replicate. A total of 3 technical and 3 biological replicates were performed. Bars represent the mean of all replicates with the standard deviation. The Mann-Whitney U Test was used to determine statistical significance, where *** is p<0.0005, ** is p<0.005, * is p<0.05, and p>0.05 is no significance.

### Urease promotes dissemination during CAUTI

To investigate the role of urease in a CAUTI, we first assessed potential strain differences between JE2 and UP 129. Bladders, catheters, kidneys, and spleens were assessed at 1-,4-, and 7-days post infection (dpi) for bacterial load and inflammation (**Figure 6, S9**). Notably, at most time points, there were no significant differences between bacterial burden or inflammation between the two strains. However, at 4-dpi UP 129 displayed significantly higher catheter colonization compared to JE2 (**Figure 6A**). Notably, this corresponded with significantly higher inflammation in bladders and significantly lower inflammation in the spleens infected with UP 129 compared to JE2 at 4-dpi (**Figure 6B, F**). These data suggest that UP 129 may have an advantage in colonizing surfaces, such as the urinary catheter. Additionally, at 7-dpi UP 129 displayed significantly lower bladder colonization compared to JE2 (**Figure S9D**).

**Figure 6:**
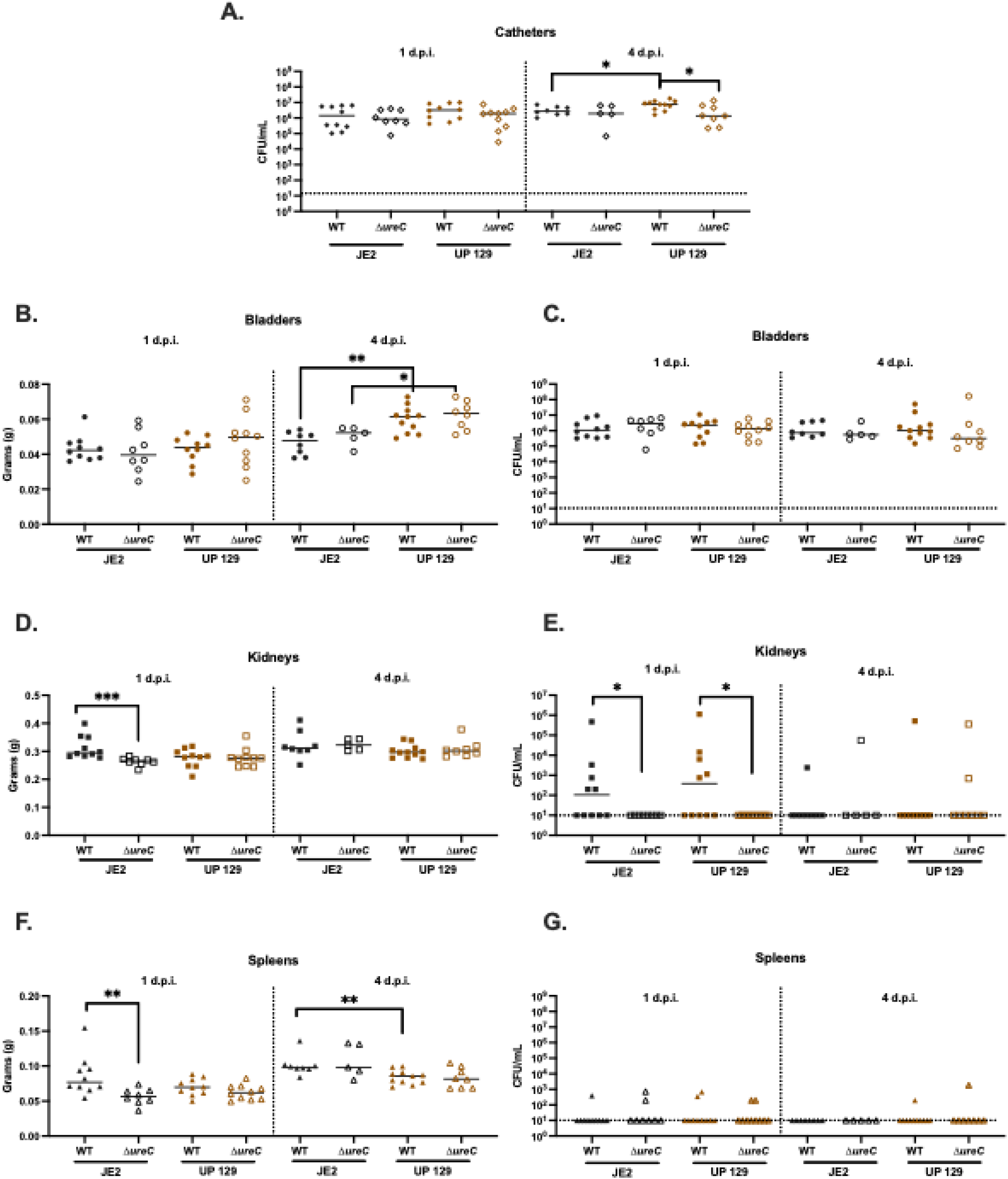
CAUTI mouse model of S. aureus wild type and ΔureC strains. Mice were infected with either JE2, JE2 ΔureC, UP 129, or UP 129 ΔureC for 1-, and 4- dpi. Colony forming units (CFUs) were obtained from the catheter (A), bladder (C), kidneys (E), and spleen (G). The weights of the bladders (B), kidneys (D), and spleens (F) were also measured at each time point. Each symbol represents one catheter, bladder, kidney, and spleen from an individual mouse, respectively. Closed black symbols indicate mice infected with JE2 and open black symbols indicate mice infected with JE2 ΔureC. Closed brown symbols indicate mice infected with UP 129 and open brown symbols indicate mice infected with UP 129 ΔureC. The Mann-Whitney U Test was used to determine statistical significance between groups, where *** is p<0.0005, ** is p<0.005, * is p<0.05, and p>0.05 is no significance. Solid lines indicate the median.

Notably, the increased inflammation within the bladder of UP 129 at 4-dpi, may explain the lower burden within the bladder compared to JE2 at 7-dpi (**Figure 6B, S9D**).

Next, to determine the contribution of urease to CAUTI, the *ureC* mutant strains were assessed in at 1-, 4- and 7-dpi. At most timepoints, there was no significant difference between organ colonization or inflammation between either wild type strain or their respective *ureC* mutant (**Figure 6, S9**). However, UP 129 *ΔureC* displayed significantly lower bacterial colonization on the catheter compared to UP 129 at 4dpi, which contrasted with JE2 (**Figure 6A**). This difference may be due to the colonization differences on the surface of the catheter between the wild type JE2 and UP 129 strains discussed above. Impactfully, at 1 dpi the *ΔureC* mutant of both background strains did not disseminate to the kidney, resulting in significantly lower bacterial loads compared to their respective parent strains (**Figure 6E**). This phenotype was ablated at 4- and 7- dpi as the mice begin to control the disseminated infection (**Figure 6E-G, S9H**). There was also an observed decrease in kidney and spleen inflammation in the JE2 *ΔureC* strain compared to the parent strain that was not observed in the UP 129 background at early time points (**Figure 6D, 6F**). Together these results suggests that urease plays a role in promoting spread from the bladder to the kidneys at early time points.

## DISCUSSION

CAUTI are the most common hospital-associated infection in the United States, and incidences of infection are increasing^8,9,11^. As the global population ages, many individuals will enter nursing homes, where the urinary catheter prevalence is higher than the general population (up to 7.3%)^53,54^. Despite *S. aureus’s* prevalence in the catheterized bladder and the severity following symptomatic infection, the uropathogen has been largely understudied. While *S. aureus* encodes numerous well studied virulence determinants, factors specifically important for CAUTI remain largely understudied^24^. Our previous data shows that among clinically collected urinary catheter-associated isolates, temporal increases in urease activity are detected, implicating this virulence factor as a contributor to CAUTI^32^. While some mechanistic studies have investigated *S. aureus* urease in the healthy urinary tract^49,55,56^, there have been no studies of how urease contributes to *S. aureus* infection in the catheterized urinary tract.

In this study, we demonstrate that a complex regulatory network controls urease expression and activity in *S. aureus* (**Figure 7A**). This complex network of regulation may explain why *S. aureus* urease is difficult to detect in a clinical setting. The standard Christensen urea agar test only detects activity in 25-50% of clinical *S. aureus* isolates, despite reports that >90% of *S. aureus* strains encode the urease operon^57–59^. *S. aureus* must sense multiple signals within the environment to both relieve repression and activate expression of urease. If those signals are absent within the assay, only low levels of urease will be detected. Notably, this regulator network contrasts with the well- studied urease producer *P. mirabilis.* Specifically, in *P. mirabilis,* the urease operon is regulated by *ureR,* which is activated in the presence of urea, making the enzyme easy to detect *in vitro*^60^. *S. aureus* does not encode a *ureR* homolog and instead must rely on this complex regulatory pathway to control urease^32^. Here, we show that the stationary phase of growth is the most robust and consistent inducer of *S. aureus* urease.

**Figure 7:**
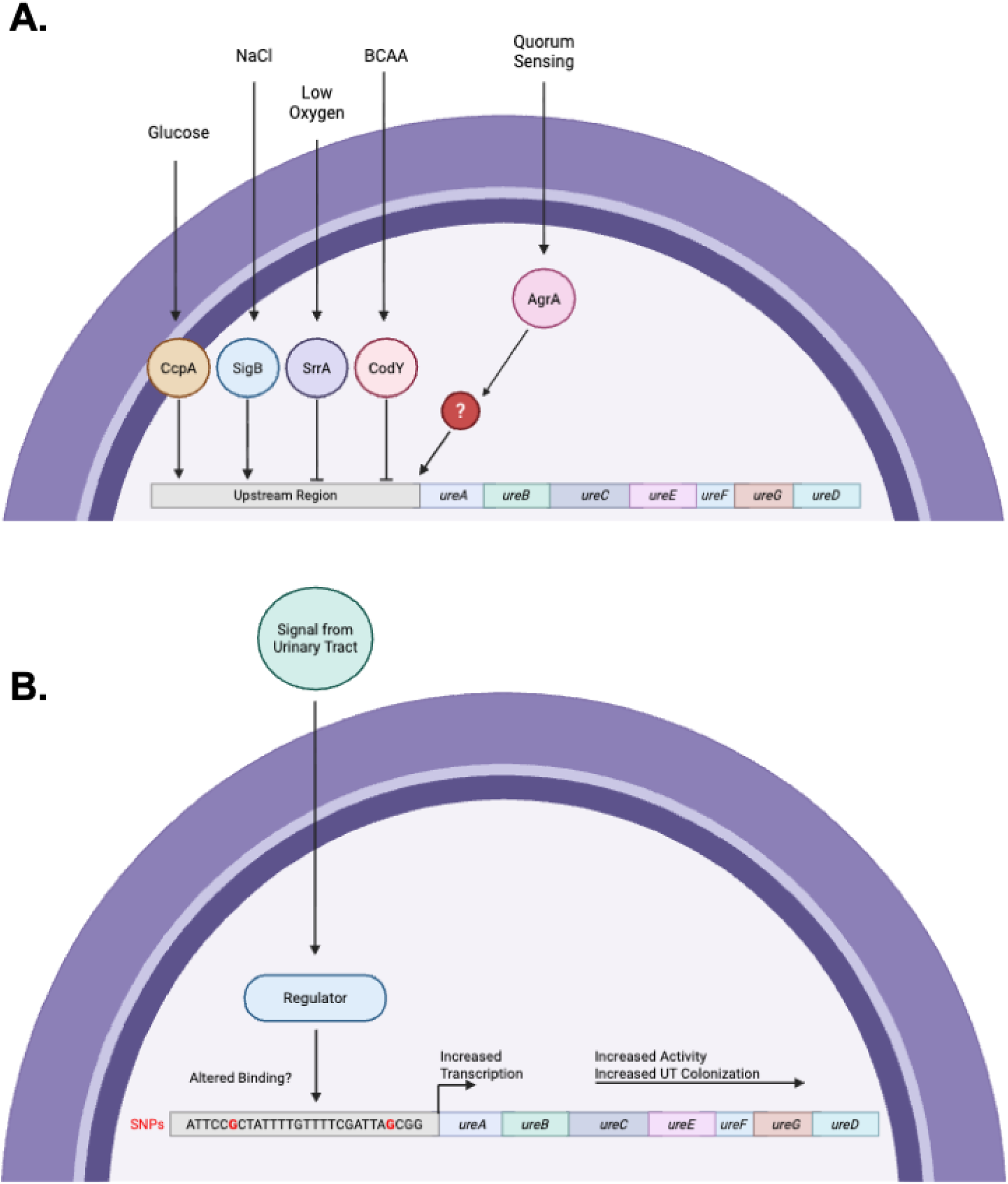
Model of urease regulation in S. aureus. A) CcpA, SigB, SrrA, CodY, and AgrA respond to their environmental signals to either activate (CcpA, SigB, AgrA), or inhibit (CodY, SrrA) urease expression and activity. CcpA, SigB, SrrA, and CodY are predicted to either promote or inhibit urease expression by binding to the upstream region of the urease operon to either recruit or block RNA Polymerase II from transcribing the operon. AgrA is predicted to act through an unknown regulator to promote urease expression. B) SNPs observed in the most common promoter region among clinically collected urinary catheter-associated isolates. SNPs are predicted to influence urease expression by altering the binding of urease regulators to the promoter. SNPs in the upstream region are sufficient to increase urease transcription, which in turn is expected to increase urease activity, and ultimately, support colonization of S. aureus in the urinary tract.

Impactfully, Christensen urea agar tests are performed with logarithmic phase cultures in media that has little to none of the activators of *S. aureus* urease in its composition. Additionally, we show that environmental conditions that mimic the urinary tract impact urease expression. Specifically, compared to standard lab media (BHI), media that mimic the urinary tract significantly increases urease expression. Notably, the phenotypes displayed following growth in AUM often mimicked the phenotypes observed in HU, yet were more consistent. The inconsistency observed for the phenotypes in HU may be due to the limited nutrients and high variability of urine composition between individuals^61^. Specifically, over 3000 metabolites have been identified in HU and the composition differs vastly between individuals and within the same individual over the course of the day^61,62^. Thus, a diversity of signals and nutrients that microbes living in the urinary tract respond to and use for survival fluctuates widely. Together, these findings may explain why *S. aureus* is labeled “a weak urease producer” in the clinic compared to *P. mirabilis*, as their mechanisms of regulation differ substantially.

Previous studies investigating the regulation of *S. aureus* urease have shown that CodY, CcpA, and Agr contribute to urease expression^27,39^. In this study, we confirmed these findings and show that two additional regulators – SigB and SrrA – also contribute to urease regulation (**Figure 7A**). Additionally, these data demonstrate that urease expression in these mutant background strains is not impacted by growth.

Specifically, strains demonstrated similar expression patterns regardless of how expression was analyzed (e.g. calculated fluorescence per cell vs fluorescence normalized to the non-fluorescent background strain). We also demonstrate that many of the signals these regulators respond to, including sugar, salt, BCAA, oxygen, and antibiotics, impact the expression of urease. Importantly, varying levels of these signals, including sugar, salt, BCAA, and oxygen, are commonly observed in the urinary tract^41–46^. Furthermore, several of the signals function through the expected regulator, including CcpA and SigB (**Figure 7A**). However, the signals that impacted urease independent of the expected regulators suggest that alternative regulation pathways are contributing to urease expression in response to these signals. Challengingly, the regulons of CodY, CcpA, SigB, Agr, and SrrA are known to display crosstalk between these regulators ^24,63–68^, which complicates the regulation of urease^23,60–65^. Specifically, CodY, CcpA, SigB, and SrrA have all been shown to regulate the expression of the Agr quorum sensing system ^24,63,64,67,68^. As such, any perturbation of these regulons may also impact Agr and subsequently urease. Additionally, SrrAB is not the only regulator that is essential for nitrous oxide resistance, as CodY is also essential for this response specifically during stationary phase^65^. This crosstalk may explain the changes in urease expression observed in the presence of nitrous oxide in the *srrA* mutant background.

These complexities necessitate further mechanistic studies to define the regulatory pathways that are directly and indirectly impacting urease expression and activity.

In previous work, we found that *S. aureus* strains that persist asymptomatically in the urinary tract display temporal increases in urease activity^12,32^. To determine whether genomic changes observed within the promoter region of these clinical isolates impact expression or activity, urease expression driven by the JE2 promoter (pUre1) or the most common promoter among the clinical isolates (pUre2) was characterized^12,32^.

While these promoters only differed by two SNPs, those genomic changes were sufficient for altering urease expression in the representative clinical isolate UP 129 background (**Figure 7B**). Interestingly, there was no difference between expression of the two promoter sequences in the JE2 background. This suggests that despite conservation across most of the urease operon and regulon, additional genomic changes, such as the missense mutations in SigB, may also play a role in regulating urease among clinical isolates. However, CC5 and CC8 are different lineages and encode diverse accessory genomes. Therefore, multiple other factors, such as other proteins in the urease regulatory pathway or changes in nickel acquisition, may play a role in influencing urease expression or activity. Future studies using allelic swaps in the JE2 and UP 129 background strains will provide additional support for this hypothesis.

One important facet of *S. aureus* CAUTI is biofilm formation. Many studies analyzing the surface of human catheters demonstrate that *S. aureus* forms complex biofilm structures on the surface of the catheter^10,13,15,18,51,69^. In this study, we show that our initial findings aligned with previous work, demonstrating that urease is not essential for biofilm formation in rich media^49^. However, by altering the conditions to more accurately reflect the catheterized bladder environment^10,51,52,70–72^, we demonstrate that urease promotes biofilm, regardless of strain background. These data also correlate with the finding that the urease operon is upregulated in biofilm cultures^48^. Furthermore, these findings emphasize the importance of urease in promoting *S. aureus* colonization in catheterized urinary tract-like conditions.

In contrast to previous work in a non-catheterized UTI model, which showed that urease did not significantly impact the pathogenesis of *S. aureus* during infection^49^, this study implicates urease as an important virulence factor during *S. aureus* CAUTI. Notably, it has been established that catheterization promotes robust, persistent *S. aureus* colonization in the urinary tract by initiating release of host factors that are bound by *S. aureus* adhesins to promote colonization^52^. Our data showed that urease was indeed important for catheter colonization, further emphasizing the importance of using models that mimic disease states. Furthermore, UP 129, a high urease producing strain, had significantly higher colonization on the surface of the catheter at 4-dpi compared to JE2. This finding correlates with our biofilm data, which shows the importance of urease in forming biofilms in a catheterized bladder-like environment.

These results may also explain the observed lower JE2 colonization on the surface of the catheter in our CAUTI model, which is a lower urease producer compared to UP 129. Impactfully, our mouse model demonstrates that urease is essential for early dissemination to the kidney, which may explain why *S. aureus* often increases the risk of dissemination to the upper urinary tract and subsequently the bloodstream^18,19^.

Overall, our findings not only demonstrate that environmental conditions play a crucial role in regulating urease, but also that strain differences can contribute to differences in expression and activity. Specifically, among clinical urinary catheter-associated isolates that are adapted to the catheterized bladder environment, urease expression and activity are higher when compared to an historical isolate. This study emphasizes the importance of using clinical isolates in understanding pathogenesis, as the phenotypes can differ between these isolates and historical strains. Future work will aim to further elucidate the mechanism of *S. aureus* urease regulators and will aim to determine which genomic changes have the greatest impact on urease expression and activity phenotypes. Deeper insight into the mechanisms that promote adaption to the bladder may support therapeutic discoveries that target *S. aureus* CAUTI, which will ultimately improve the quality of life of individuals who have long-term urinary catheters.

## Materials and Methods

### Bacterial Strains and Growth Conditions

All strains used in this study are listed in **Table 1**. JE2 and all transposon mutants used in this study were obtained from BEI Resources and the *S. aureus* clinical isolate UP 129 was collected as part of a previously published cohort of urology outpatients with long-term indwelling urinary catheters^11,12^. Brain Heart Infusion (BHI) broth (Becton Dickinson and Company, REF #237500) and agar (Fisher, CAS #9002-18-0) was used to grow and maintain all *S. aureus* cultures for experiments. Transposon mutants were selected for and maintained with BHI or tryptic soy broth (TSB) (Sigma-Adrich, REF #22092-500G) supplemented with 50 ug/mL of erythromycin (Acros Organics, CAS #114-07-8). The pJB38 *ΔureC* plasmid was maintained in *Escherichia coli* DH5α in LB Broth (Fisher, REF #BP1426-500) supplemented with 50 ug/mL of ampicillin (Fisher, CAS# 69-52-3) at 37°C and maintained in *S. aureus* in BHI supplemented with 50 ug/mL of chloramphenicol (Sigma Aldrich, REF #C0378-25G) at 30°C. Once homologous recombination was complete, the *ΔureC* strains were maintained in BHI with no antibiotic. pUre plasmids were generated in *E. coli* DH5α in LB supplemented with 50 ug/mL ampicillin and maintained in *S. aureus* in BHI supplemented with 50 ug/mL of erythromycin and/or chloramphenicol. Urease broth was made as previously described^32^. Briefly, 1 g of peptone (BD Biosciences, REF #211921), 1 g glucose (Sigma-Aldrich, REF #G8270-1KG), 2 g potassium phosphate monobasic (Sigma- Adrich, REF #P9791-100G), 5 g sodium chloride (Fisher, CAS #7647-14-5), and 0.012g Phenol red (Sigma-Aldrich, REF #114529-5G) were resuspended in 1 L of water. Media was sterilized and was supplemented with filter sterilized 40% urea solution (Fisher, REF#BP169-500). Artificial urine media (AUM) was adapted based on previously published artificial urine recipes. Briefly, AUM was prepared as previously described^61^, with the exception that an additional 0.909 mM tryptone (Becton Dickenson and Company, REF #211705), 0.833 mM glucose (Fisher, CAS #50-99-7), 82 nM biotin (Thermo, Cat #230090010), and 0.17% yeast nitrogen base (YNB) without amino acids and ammonium sulfate (Becton Dickenson and Company, REF #233520) was added.

**Table 1.** Strains used in this study.

| Denotation in this Paper | Isolation Site | Reference | Denotation in Reference |
| --- | --- | --- | --- |
| JE2 | Skin | <sup>73</sup> | JE2 |
| JE2 $\Delta ureC$ | Skin | This study | JE2 $\Delta ureC$ |
| JE2 *codY::tn* | Skin | <sup>74</sup> | JE2 *codY::tn* |
| JE2 *ccpA::tn* | Skin | <sup>74</sup> | JE2 *ccpA::tn* |
| JE2 *sigB::tn* | Skin | <sup>74</sup> | JE2 *sigB::tn* |
| JE2 *agrB::tn* | Skin | <sup>74</sup> | JE2 *agrB::tn* |
| JE2 *saeR::tn* | Skin | <sup>74</sup> | JE2 *saeR::tn* |
| JE2 *srrA::tn* | Skin | <sup>74</sup> | JE2 *srrA::tn* |
| RN4220 | Cloning Strain | <sup>75</sup> | RN4220 |
| UP 129 | Urinary Catheter | <sup>11</sup> | Urology outpatient<br>129 |
| UP 129 $\Delta ureC$ | Urinary Catheter | This study | UP 129 $\Delta ureC$ |

To adjust the buffering capacity changes associated with the addition of YNB, the amount of sodium phosphate monobasic (Mallinckrodt Pharmaceuticals, CAS #10049- 21-5) was adjusted to a total of 18.667 mM.

### Generation of Chloramphenicol Resistant GFP Transcriptional Fusion Plasmid

GFP transcriptional fusion plasmids compatible with the transposon mutant strains (erythromycin cassette) were generated by inserting a chloramphenicol resistance cassette to the pCM11 plasmid, as previously described^76^. Briefly, the pGFP chloro F and pGFP chloro R primers flanked with XbaI and PstI restriction sites (**Table 2**) were used to amplify the chloramphenicol cassette from the pJB38 plasmid^77^. The PCR product was cloned directly into the pCR4-Blunt TOPO plasmid using the Zero Blunt TOPO PCR Cloning Kit (Thermo Fisher, Cat #450031) according to the manufacturer’s protocol. The plasmid was then transformed into *E. coli* DH5α. The pCR4-Blunt TOPO- chloramphenicol cassette plasmid and the pCM11 plasmid were then digested using the XbaI and PstI restriction enzymes. Both fragments were gel purified and ligated with T4 DNA ligase. The resulting plasmid (pGFPchloro/erm) was then transformed into DH5α and was confirmed via sequencing (**Supplemental File 1**). The pGFPchloro/erm plasmid was then used as a backbone for GFP transcriptional fusions of the upstream region of the urease operon.

**Table 2.**
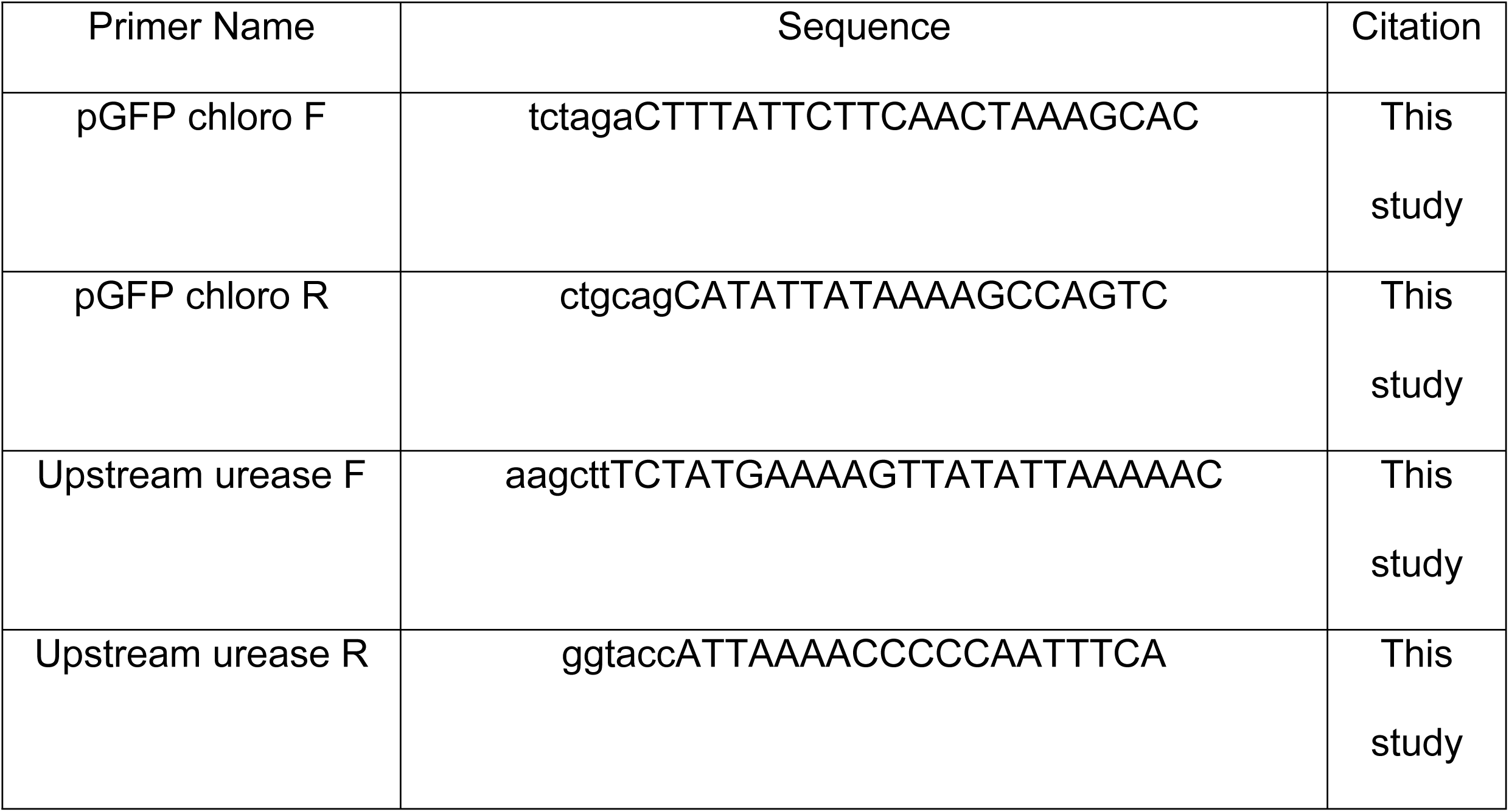
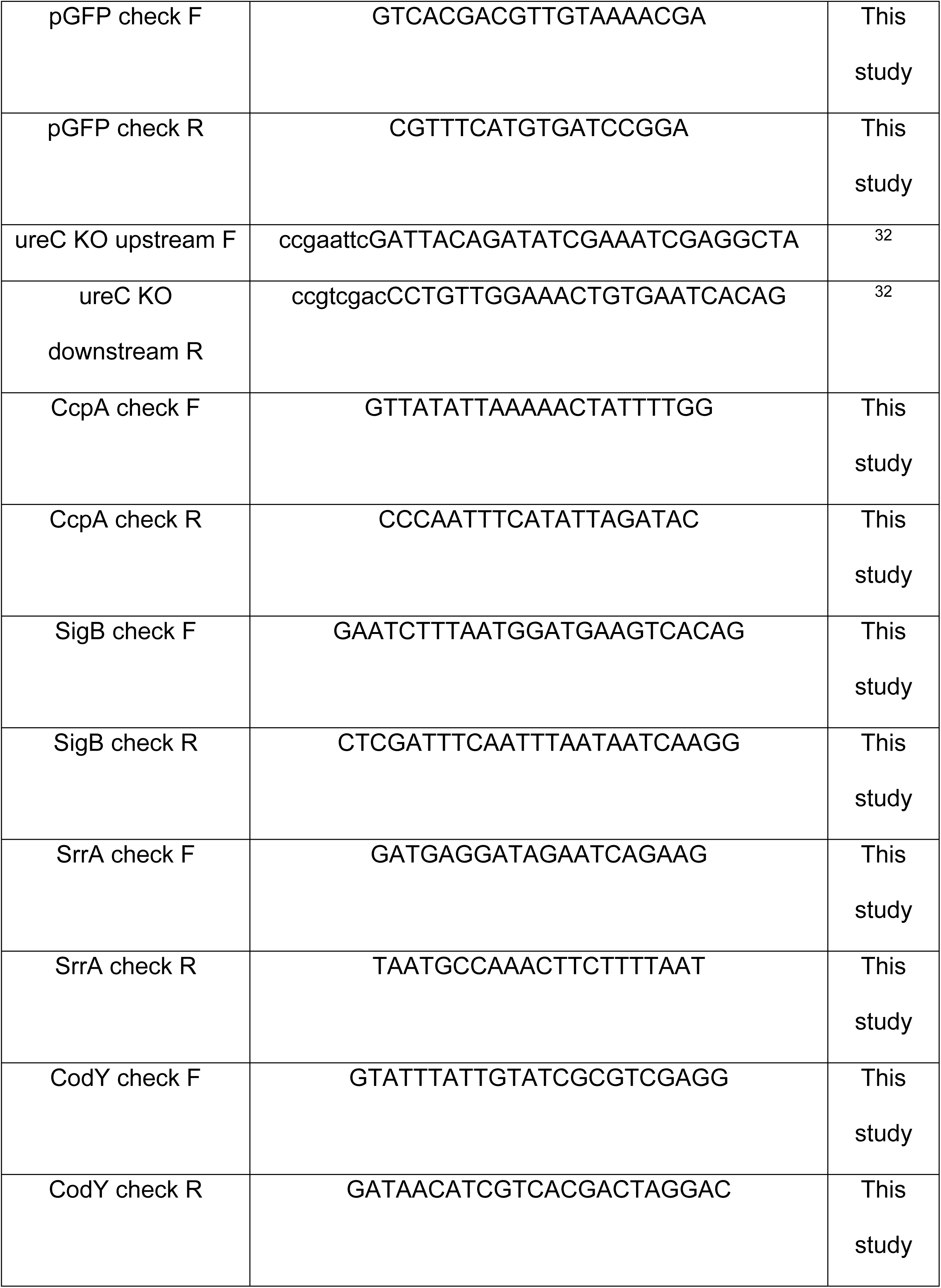
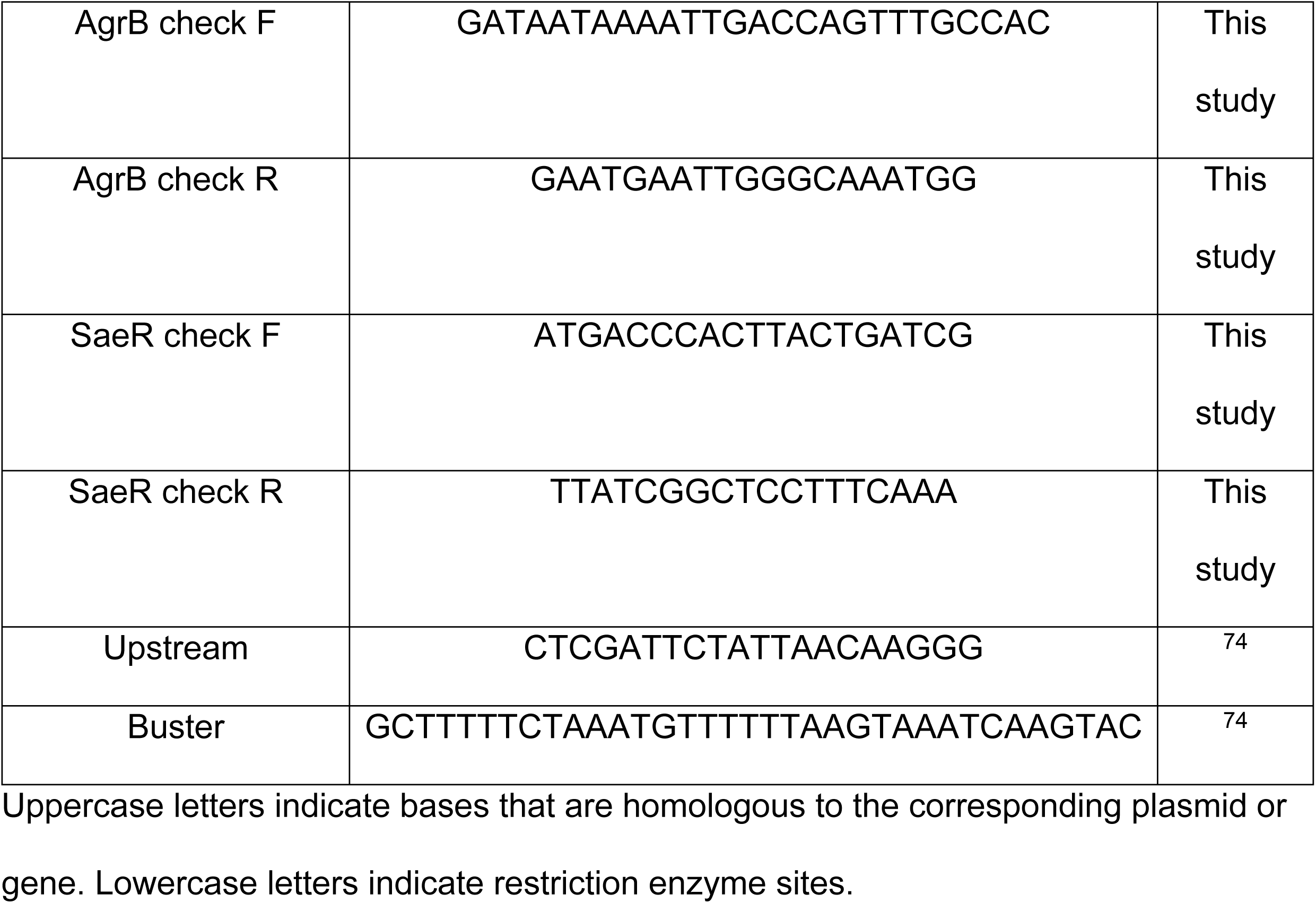
Primers used in this study.

Uppercase letters indicate bases that are homologous to the corresponding plasmid or gene. Lowercase letters indicate restriction enzyme sites.

### Generation of Urease Transcriptional GFP Fusion Constructs

GFP transcriptional fusions to the upstream region of the urease operon were generated using the pGFPchloro/erm plasmid, as previously described^78^. Briefly, the upstream region of urease (200bp) was amplified using the forward (upstream urease F) and reverse (upstream urease R) primers flanked with HindIII and KpnI restriction sites (**Table 2**). The PCR product was cloned directly into the pCR4-Blunt TOPO plasmid, using the Zero Blunt TOPO PCR Cloning Kit and transformed into DH5α. The plasmid was then isolated and digested using HindIII and KpnI restriction enzymes.

Following gel purification, the upstream region fragment was ligated into pGFPchloro/erm, which was similarly digested with HindIII and KpnI. The resulting plasmids (pUre) were confirmed via PCR and sequencing (**Supplemental File 1**). pUre plasmids were then transformed into the *S. aureus* lab strain RN4220 and subsequently transduced via phage 11 into the JE2, as previously described. The pUre plasmids were introduced into the UP 129 strain via electroporation, as previously described^32^. Final plasmids were confirmed via PCR (**Figure 7C**).

### GFP Transcriptional Fusion Expression Assays

Transcriptional fusion expression assays were performed based on previous work^32,78^. Overnight cultures were grown at 37°C in BHI. For analysis of logarithmic phase cultures, strains were diluted 1:100 from overnight culture in various growth media, including BHI, AUM, pooled female HU (see Human urine collection section below), a 1:1 ratio of pooled female HU and saline, and urease broth^32^ without phenol red. For analysis of stationary phase cultures, 1 mL of culture was washed in 1X PBS and then resuspended in various growth media. Cultures were transferred to a clear bottom, black walled 96-well plate and OD_600_ and fluorescence were measured every hour for a total of 18 hours in the Synergy H1 Biotek microtiter plate reader (Biotek, Vermont).

GFP excitation wavelength was 488 nm and the emission was collected at 515 nm, as previously described^79^. Fluorescence of each strain was normalized by subtracting the fluorescence from the background strain (no GFP reporter plasmid plasmid) from each strain that contained the GFP reporter plasmid. To determine fluorescence per cell, the ratio of GFP over the OD_600_ was measured. This ratio was normalized by subtracting the ratio from the background strain (no GFP reporter plasmid) from each strain that contained the GFP reporter plasmid.

### Ethics Statement

For all patient samples obtained in this study, formal written consent was obtained. All samples were obtained in accordance with protocol IRB# HSC-MS-21-0849 (human urine collection) or protocol #IRB HSC-MS-23-0163 (human plasma collection) and were approved by the Committee for the Protection of Human Subjects (CPHS) at the University of Texas Health Science Center at Houston – UTHealth and performed in accordance with the 1964 Helsinki Declaration and its later amendments.

### Human Urine Collection

Human urine was sourced from the Urine BioBank for Urinary Tract Infection Studies, in accordance with protocol IRB# HSC-MS-21-0849 approved by the CPHS at the University of Texas Health Science Center at Houston – UTHealth. Briefly, donors were consented and enrolled prior to donation. Urine from at least two healthy female donors, aged 18+ were collected and pooled. Donors were not taking antibiotics, nor currently menstruating, pregnant, and or experiencing symptoms of a UTI. Urine was sterilized with a 0.22 um filter (Thermo, Cat #595-3320) and stored at 4°C until use. A one to one ratio of HU and saline (HU:S) solution was created by combining one volume of HU to one volume of 0.9% saline solution.

### Identification of Consensus Regulator Binding Sites

Consensus regulator binding sites within the regulatory region of the urease operon were identified using the program Artemis (Sanger, version 18.1.0) to analyze the 200- base pair sequence upstream of the urease operon. 171 *S. aureus* isolates were previously isolated from long-term catheterized individuals and their genomes were sequenced (**Supplementary File 1**)^9,11,12^. The sequence of the *S. aureus* historic strain JE2 was obtained from NCBI (**Supplementary File 1**). The urease operon and upstream region were identified in each genome and consensus binding sequences of predicted regulators were identified based on the reported binding consensus sequences in the literature^33–37,80^. Alignment of the 200-base pair region upstream of the urease operon for the 171 urinary catheter-associated isolates^11,12^ and JE2 was created using the DECIPHER software package^81^ in R Studio (Posit Software, version 2023.06.0+421).

### Transposon Mutant Generation

Mutations in predicted urease regulators were reconstructed in the JE2 wildtype strain background using the Nebraska Transposon Mutant Library (NTML), as previously described^32^. Briefly, phage transduction, using phage 11, was used to generate lysates of each transposon mutant regulator. The resulting phage lysate was used to backcross each transposon mutant into the JE2 strain background and were selected for using TSA supplemented with 20 mM sodium citrate (CAS #6132-04-3) and 50 ug/mL erythromycin. Transposon mutants were confirmed via PCR (**Figure S3A**) using the forward and reverse regulator primers and the upstream or Buster primers listed in **Table 2**.

### Generation of *ureC* Clean Deletion Strains

The *ΔureC* strains were generated as previously described^32^. Briefly, the previously generated pJB38 *ΔureC* plasmid^32^ was isolated using a plasmid miniprep kit (Promega, REF# A1330) following the manufacturers protocol, with the exception that 10 ug/mL of lysostaphin was used to lyse *S. aureus* cells (Sigma Aldrich, REF# L7386-5MG).

Electrocompetent cells of JE2 and UP 129 were generated as previously described^32^. Briefly, overnight cultures were subcultured into 50 mL of B2 medium (1% casein hydrolysate (Oxoid, REF #LP0041), 2.5% yeast extract (Fisher, REF #BP1422-500), 0.5% glucose (Sigma Aldrich, REF #G8270-1KG), 2.5% NaCl (Fisher, REF #S671-500), 0.1% K2HPO4 (Sigma Aldrich, REF #P8281-100G)) at an OD_600_ of 0.25. Cells were grown at 37°C for ∼2 hours until OD_600_ 0.4. Cells were then washed in 50 and 12.5 mL of cold, sterile water followed by 7 mL of cold 10% glycerol. Finally, cells were resuspended in 200 uL of cold 10% glycerol and were stored at -80°C until use.

Electrocompetent cells were transformed with purified pJB38 *ΔureC* plasmid and transformants were selected for at 30°C on chloramphenicol plates. Co-integrates were selected for at 42°C overnight on chloramphenicol plates and resulting colonies were grown in TSB without antibiotic at 30°C. Recombinants were counter-selected for with 100 ng/mL of anhydrotetracycline hydrochloride (Fisher, Cat #AAJ66688MA). Colonies were screened for both urease activity and chloramphenicol sensitivity. The deletion of the *ureC* gene was further confirmed via PCR (**Figure S9A**).

### Urease Activity Assays

Urease activity was measured as previously described^32^. Briefly, bacterial strains were grown in BHI for 18 hours. Cultures were washed in 1X PBS and resuspended in urease broth, either with or without urea. Each strain was transferred to a U-bottom 96- well plate (Greiner, Cat #650185), in triplicate, with a minimum of two biological replicates. The OD was measured at 415, 560, and 600 nm every hour for 18 hours while shaking at 37°C in the BioTek Synergy H1 microtiter plate reader. The ratio of ODs at 560 nm and 415 nm was used to calculate the color change in the media, which corresponds to the change in pH. This color change was normalized to the urea negative controls to account for any color change that was not due to urease activity.

### Human Blood Collection and Plasma Extraction

Human blood was collected in accordance with protocol #IRB HSC-MS-23-0163 approved by the CPHS at the University of Texas Health Science Center at Houston – UTHealth. Briefly, donors were consented and enrolled prior to donation. Blood was drawn by a trained medical professional using a 22-gauge blood collection needle (BD, REF 368652) holder, and a tube (Henry Schein, ref #9870790) containing heparin.

Human plasma was extracted from blood samples as previously described^82^. Briefly, the tube containing heparin and blood was mixed with equal volumes of 3% dextran in 0.9% saline and allowed to incubate statically at room temperature for 20 minutes to allow red blood cells to settle. The top plasma layer was removed, spun down for 10 minutes at 2000 rpm at 4°C, and the supernatant was aliquoted in 1.7 mL tubes for storage at - 20°C until use.

### Biofilm Assays

Biofilm biomass was measured as previously described^83^. Briefly, flat-bottom 96-well plates (Greiner, Cat #655185) were coated with 50 mM sodium bicarbonate (in vitro biofilm assay) or 20% human plasma (HP) diluted in 50 mM sodium bicarbonate overnight at 4°C (host protein supplemented biofilm assay). The next day, excess HP (or sodium bicarbonate alone) was removed, and overnight cultures were diluted to an OD_600_ of 0.2 in various media, placed in the pre-coated 96-well plate, and incubated at 37°C overnight with gentle shaking. The next day, non-adherent bacteria were removed, wells were air-dried for 30 minutes, washed with water, and stained with 0.1% crystal violet for 10 minutes. Wells were washed again to remove excess crystal violet and remaining crystal violet was solubilized in 33% acetic acid overnight at 4°C. The next day, 50 uL of each well was mixed with 50 uL of 33% acetic acid and the OD_600_ was measured. A dilution factor of 1:2 was accounted for on the OD_600_ to determine the biomass of each biofilm.

### CAUTI Mouse Model

The mouse CAUTI model was performed as previously described^52,84^. Briefly, C57BL/6 female mice (Charles River Labs) were transurethrally implanted with a 3 mm piece of polyethylene tubing (Braintree Scientific; Cat #PE-10) encased in a 4 mm piece of silicone tubing (Braintree Scientific; Cat #SIL-205) and inoculated with approximately 2 x 10^7^ CFUs of bacteria. Bacterial strains were prepared for inoculation following growth in BHI for 18 hours at 37°C, centrifugation at 10,000 rpm for 10 minutes, washing with 1X PBS, and finally resuspending in 1X PBS. After 1-, 4-, or 7-days post infection, bladders, kidneys, spleens, and catheters were harvested, organs were weighed, homogenized, and CFUs were enumerated to determine bacterial load. All animal work was approved by the Institutional Animal Care and Use Committee (IACUC) and Animal Welfare Committee (AWC) at the University of Texas Health Science Center at Houston – UTHealth (protocol # AWC-23-0049) and were performed in accordance with the protocol.

### Statistical Analysis

Figures were graphed and statistics were performed using GraphPad Prism 10.4.2. All experiments included at least 2 biological replicates each, with 3 technical replicates in each biological replicate. For mouse experiments, each mouse was individually graphed in addition to the mean of each group. To reach a power of 0.8 using an alpha value of 0.05, the Mann Whitney U test power calculations determined a minimum of 8 mice per experimental group were required for the mouse model. Statistical analysis was performed with the Mann-Whitney U test with GraphPad Prism (version 10.5.0) software to determine significant differences between the wild-type and urease mutant strains.

Transcriptional analysis graphs include the mean of each group and the standard deviation from the mean. The Mann Whitney U Test was used to compare transcription between various environmental conditions. Urease activity was graphed over time for 12 hours. The area under each curve was then determined and compared between strains using the Mann Whitney U Test.

## ACKNOWLEDGEMENTS

The *codY, ccpA, sigB, agrB, saeR,* and *srrA* transposon mutants used in this study were provided by the Network on Antimicrobial Resistance in *Staphylococcus aureus* (NARSA) for distribution through BEI Resources, NIAID, NIH: Nebraska Transposon Mutant Library (NTML) Screening Array, NR-48501.

## SUPPORTING INFORMATION

**Figure S1:**
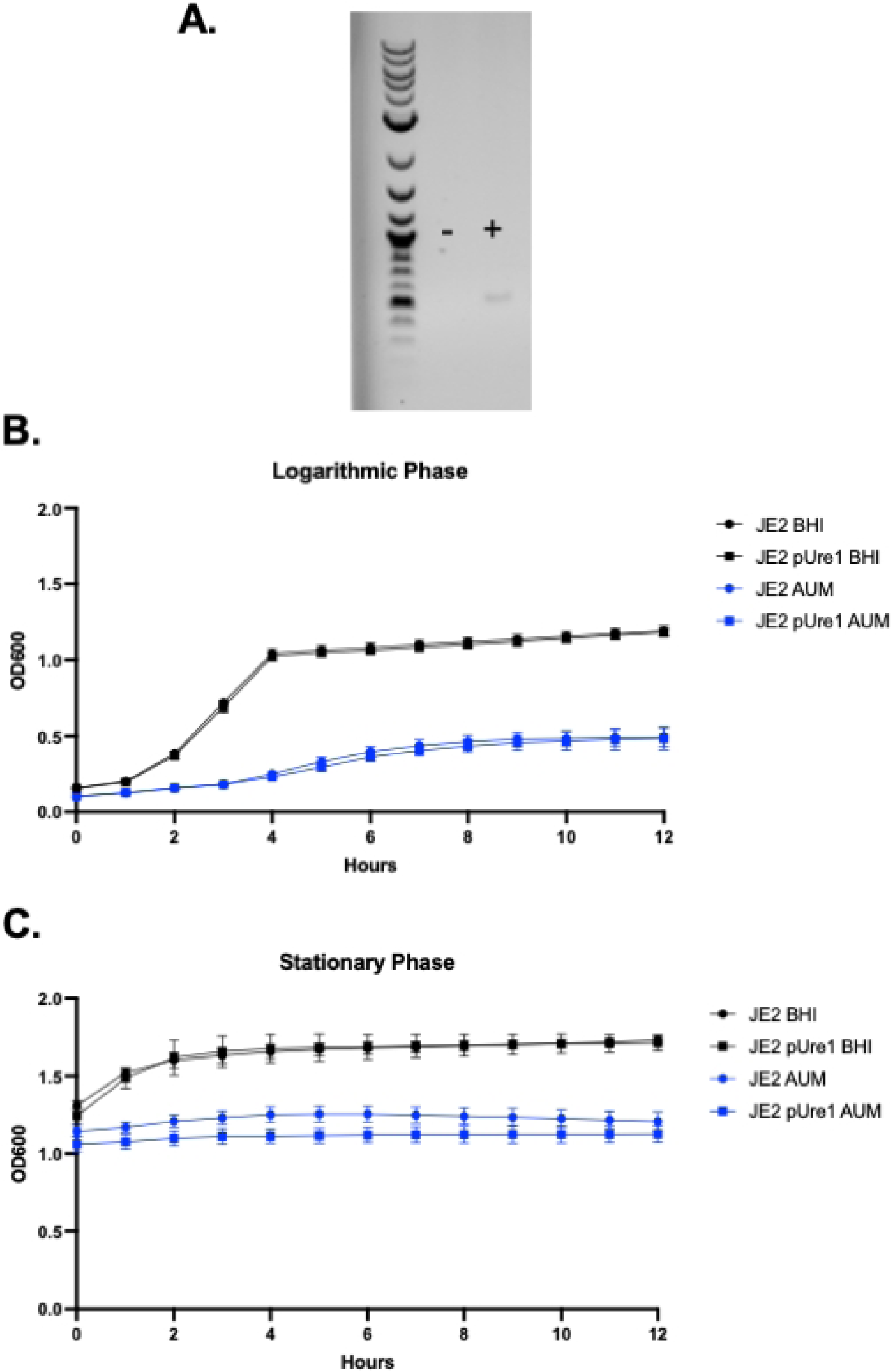
Creation and growth of JE2 pUre1 strain. **A)** Confirmation of *pUre1* plasmid in the JE2 background. Presence of the *pUre1* plasmid was confirmed with the pGFP check F and R primers found in Table 2. (-) indicates wild type JE2 DNA and (+) indicates JE2 pUre1 DNA. **B and C)** Growth of the JE2 and JE2 pUre1 strains in BHI and AUM. Cultures were started at either logarithmic phase (**B**) or stationary phase (**C**). Growth was measured via OD_600_ for a 12-hour period in either BHI (black lines) or AUM (blue lines). Dots indicate the mean of all replicates and lines indicate the standard deviation.

**Figure S2:**
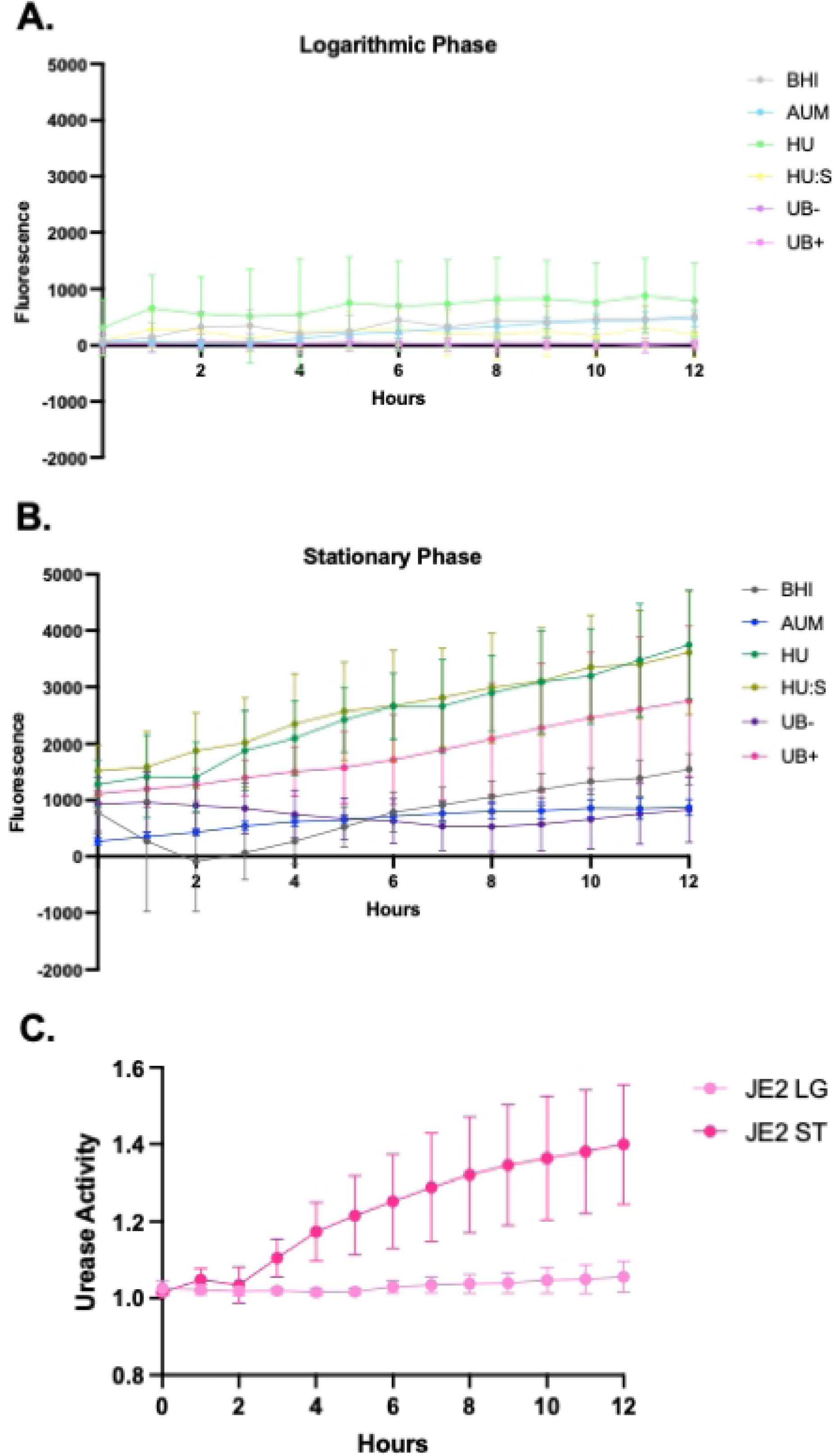
Urease expression and activity curves. Urease expression over a 12 hr time course with cells beginning in either logarithmic phase (LG) (**A**) or stationary phase (ST) (**B**) in various media conditions. Fluorescence was normalized to the fluorescence reading of the background strain that did not contain the GFP reporter plasmid. Fluorescence was measured in BHI (black lines), AUM (blue lines), female HU (green lines), HU:S (yellow lines), UB with no urea (purple lines), or UB with urea (pink lines). **C**) Relative urease activity over a 12 hr time course with cells beginning in either logarithmic phase or stationary phase.

**Figure S3:**
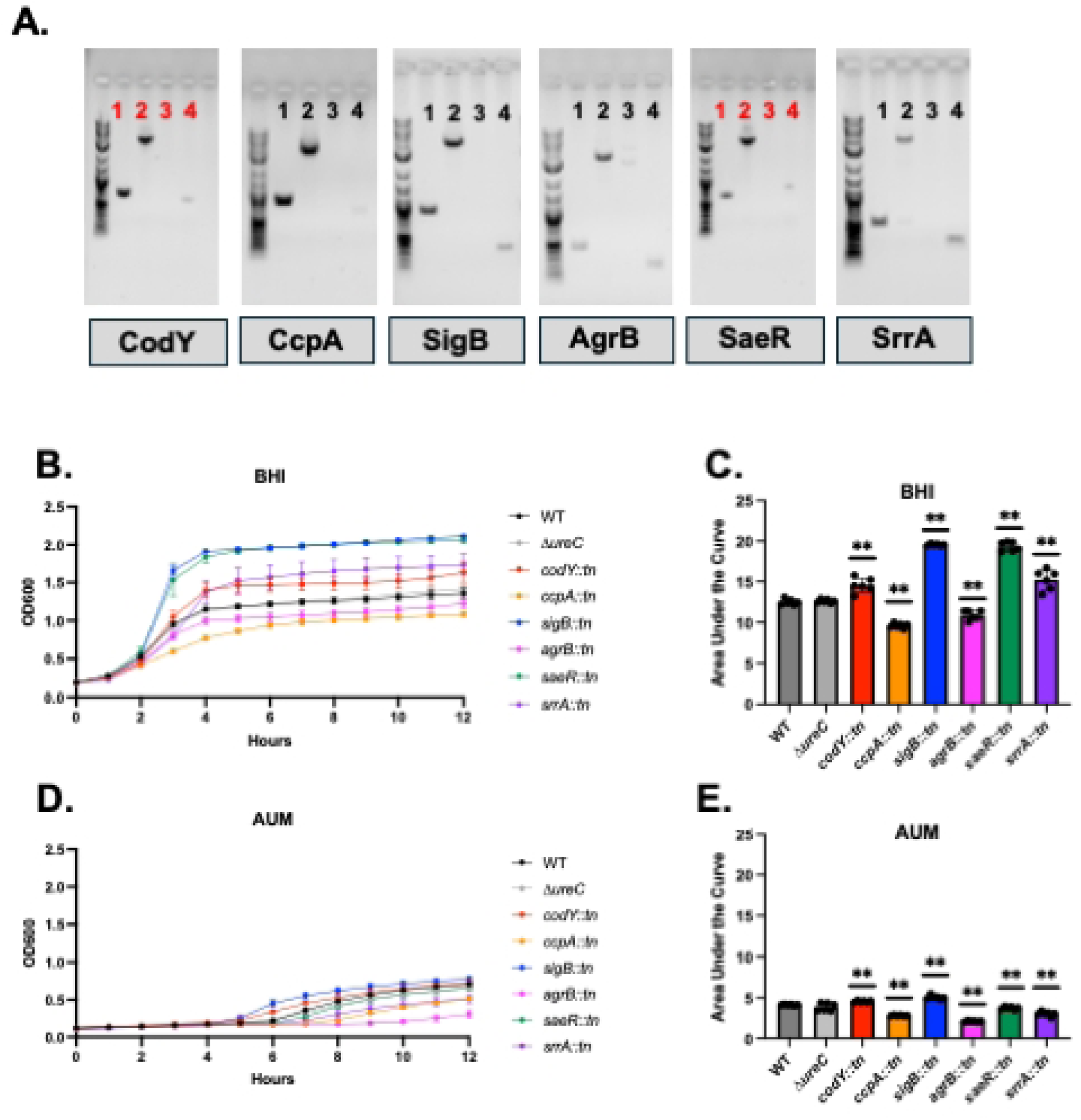
Generation and growth of transposon mutants of urease regulators. **A)** PCR confirmation of transposon mutants of *codY, ccpA, sigB, agrB, saeR,* and *srrA* in the JE2 background strain. The forward and reverse primers for each gene (Table 2) were used to amplify wild type JE2 DNA (1) and the JE2 transposon mutant DNA (2). Due to the presence of the transposon, the wild type DNA (1) had a smaller band size compared to the corresponding transposon mutant DNA (2). The forward primer of each gene and the buster primer (Table 2) were used to amplify wild type JE2 DNA (3) and the JE2 transposon mutant DNA (4). Due to the lack of a transposable element in the wild type (3), no band was amplified. The presence of the transposable element in the mutant strains allowed for binding and amplification of the buster primer (4). Growth and area under the curve analyses for transposon mutant strains in BHI (**B and C**) and AUM (**D and E**). Growth was measured via OD600 for a 12-hour period. Each dot represents a technical replicate. 2 total biological replicates were performed. Bars represent the mean of all replicates with the standard deviation. Area under the curve was calculated for each strain (**C and E**). Significance was determined using the Mann-Whitney U test.

**Figure S4:**
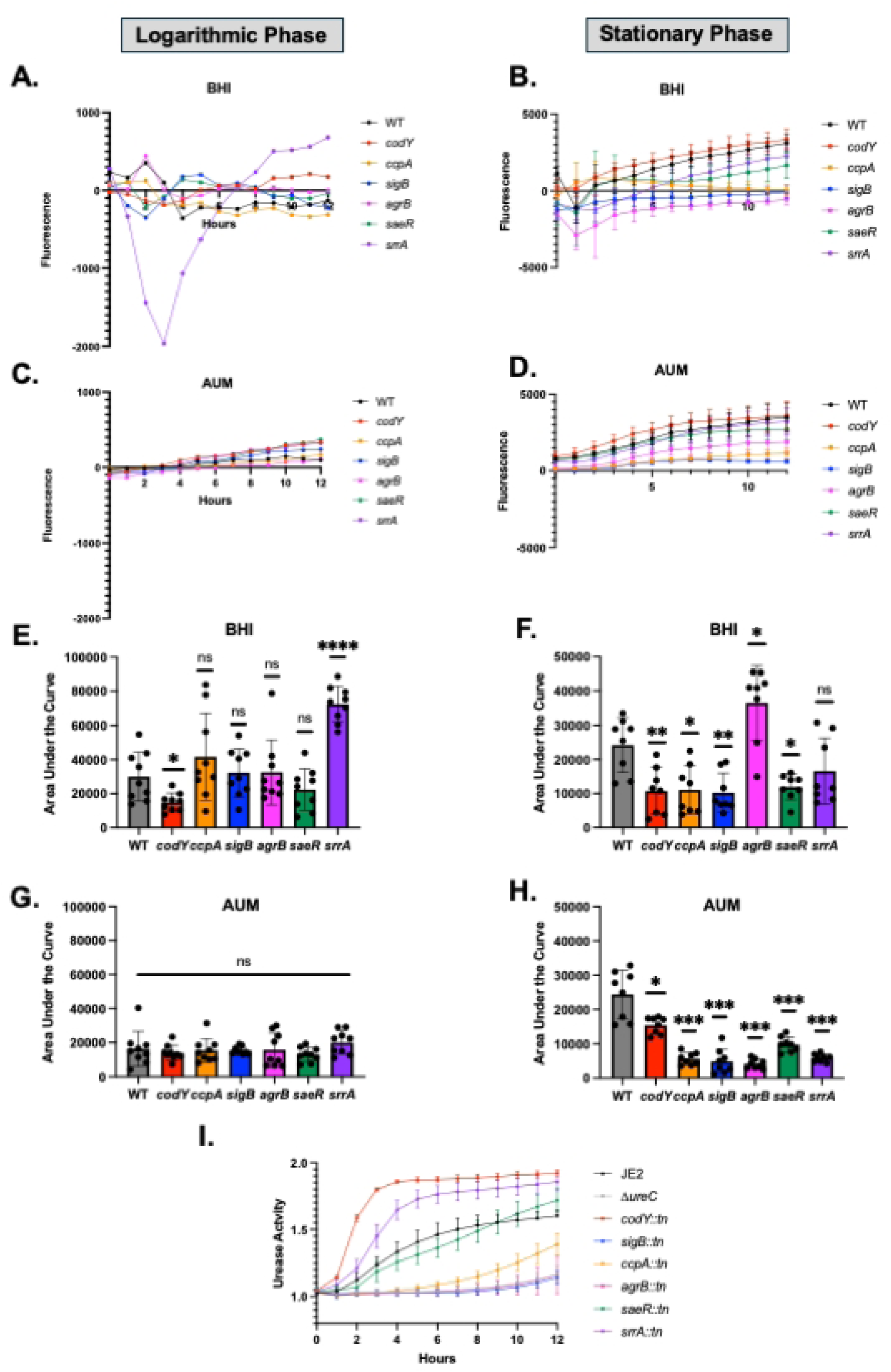
Role of predicted regulators in urease expression and activity. Urease expression over a 12 hr time course for the regulator mutant strains in either logarithmic (**A, C, E, G**) or stationary (**B, D, F, H**) growth in BHI (**A, B, E, F**) or AUM (**C, D, G, H**). Each dot represents a technical replicate. 2 total biological replicates were performed. Bars represent the mean of all replicates with the standard deviation. Area under the curve was quantified for each strain (**E-H**) and significance was determined with the Mann Whitney-U Test. **I)** Relative urease activity of regulator mutants over a 12-hour time course.

**Figure S5:**
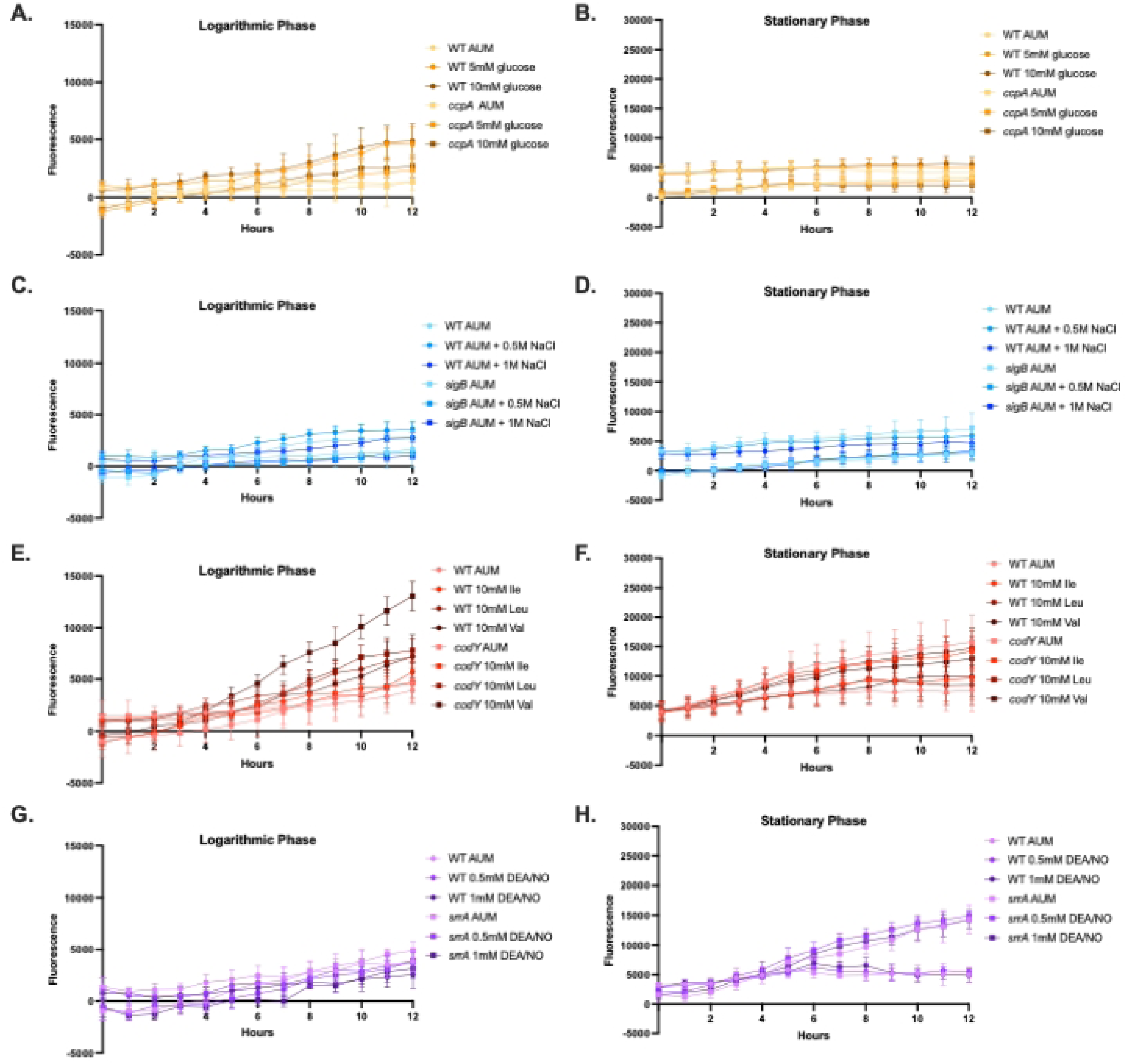
Expression curves of urease in response to various environmental stimuli. Urease expression over a 12 hr time course in the presence of environmental signals of CcpA (**A and B**), SigB (**C and D**), CodY (**E and F**), and SrrA (**G and H**). Expression was analyzed in cells in logarithmic phase (**A,C,E,G**) and stationary phase (**B,D,F,H**).

**Figure S6:**
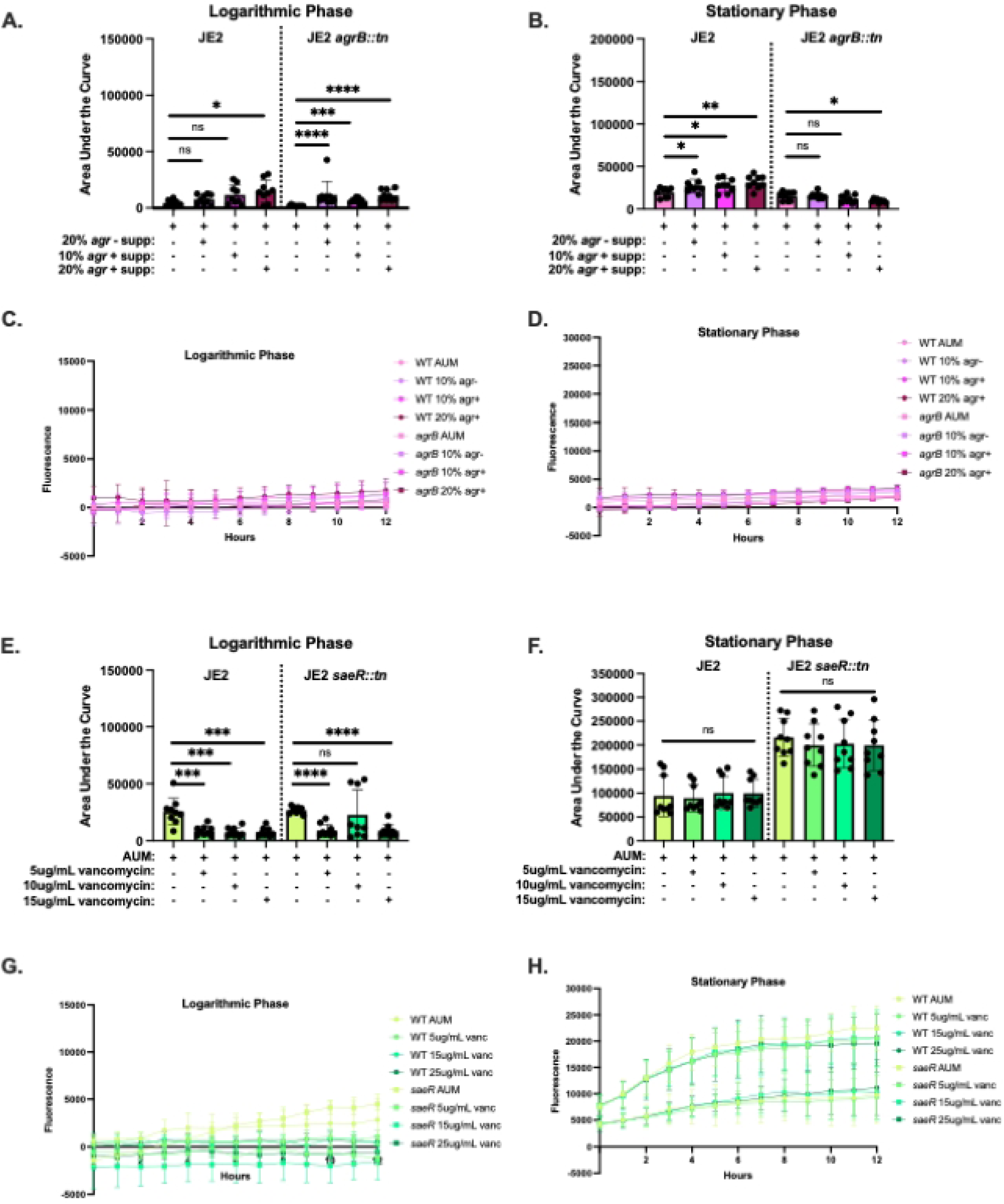
Urease expression in the presence of Agr and SaeR signals. Expression was analyzed in cells in logarithmic phase (**A,C,E,G**) and stationary phase (**B,D,F,H**) after supplementation of increasing concentrations of supernatant with or without auto inducing peptide (Agr) or vancomycin (SaeR). Each dot represents a technical replicate. 2 total biological replicates were performed. Bars represent the mean of all replicates with the standard deviation. Area under the curve analysis for AgrB signals (**M and N**) and SaeR signals (**O and P**) was analyzed and significance was determined using the Mann-Whitney U test.

**Figure S7:**
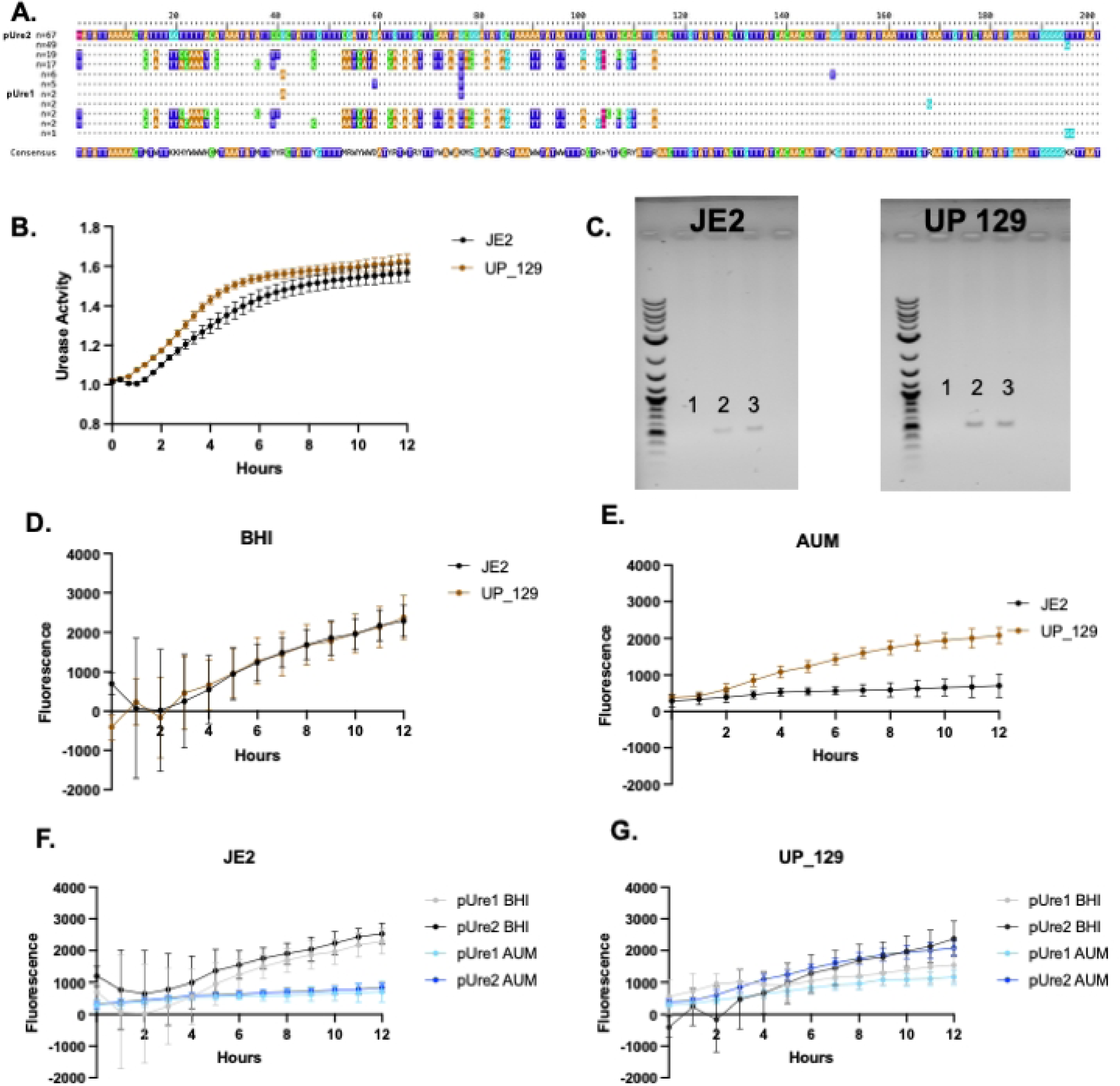
Promoter swaps of the upstream region of urease in JE2 and UP 129 **A)** Alignment of the 200 base pairs upstream of the urease promoter for JE2 and 170 urinary catheter-associated isolates. Dots indicate conservation of a nucleotide to the consensus and letters indicate a variation of nucleotides compared to the consensus. The JE2 sequence (*pUre1*) was one of the least common sequences found in these isolates, while the UP 129 sequence (*pUre2*) was the most commonly observed sequence. **B)** Relative urease activity of JE2 and UP 129 strains. Black lines indicate JE2 urease activity and brown lines indicate UP 129 urease activity. **C)** PCR confirmation of the presence of the *pUre1* and *pUre2* plasmids in JE2 and UP 129. DNA from wild type strains (1), *pUre1* strains (2), and *pUre2* strains (3) were amplified with the pGFP check primers listed in Table 2. Urease expression over a 12 hr time course of JE2 and UP 129 strains in BHI (**D**) and AUM (**E**) during the stationary phase. Black lines indicate JE2 fluorescence and brown lines indicate UP 129 fluorescence. Promoter swap expression curves for JE2 (**F**) and UP 129 (**G**). Fluorescence was measured in stationary phase cultures for 12 hours in either BHI (black lines) and AUM (blue lines).

**Figure S8:**
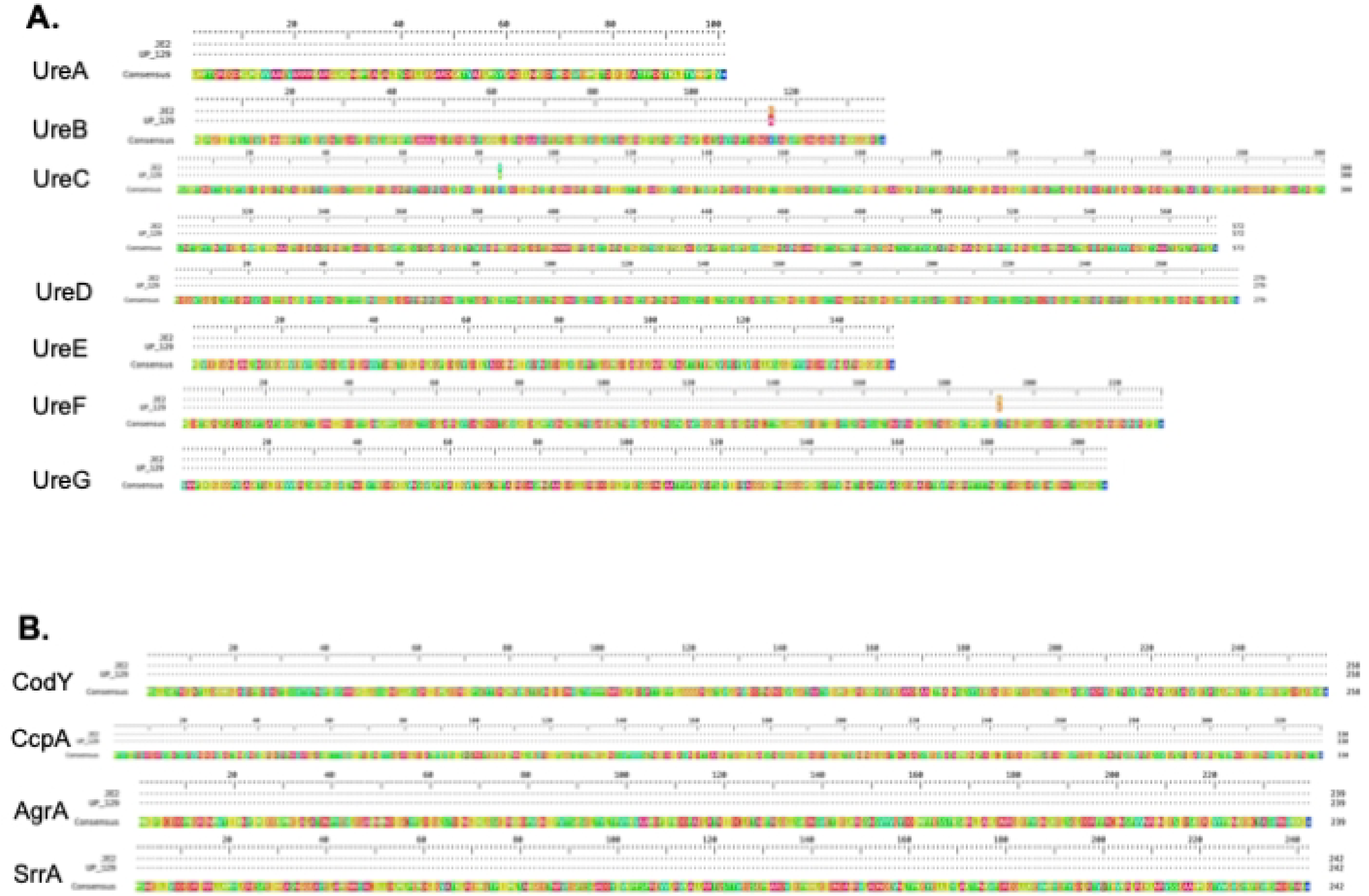
Genomic comparisons of the urease regulation pathway between JE2 and UP 129. **A)** Comparison of the gene sequences of the urease operon in JE2 to UP 129. Dots indicate conserved amino acids between JE2 and UP 129, and letters indicate SNPs resulting in missense mutations of JE2 and UP 129. **B)** Comparison of the gene sequences of CodY, CcpA, AgrA, and SrrA in JE2 and UP 129. Dots indicate conserved amino acids between JE2 and UP 129, and letters indicate SNPs resulting in missense mutations of JE2 and UP 129.

**Figure S9:**
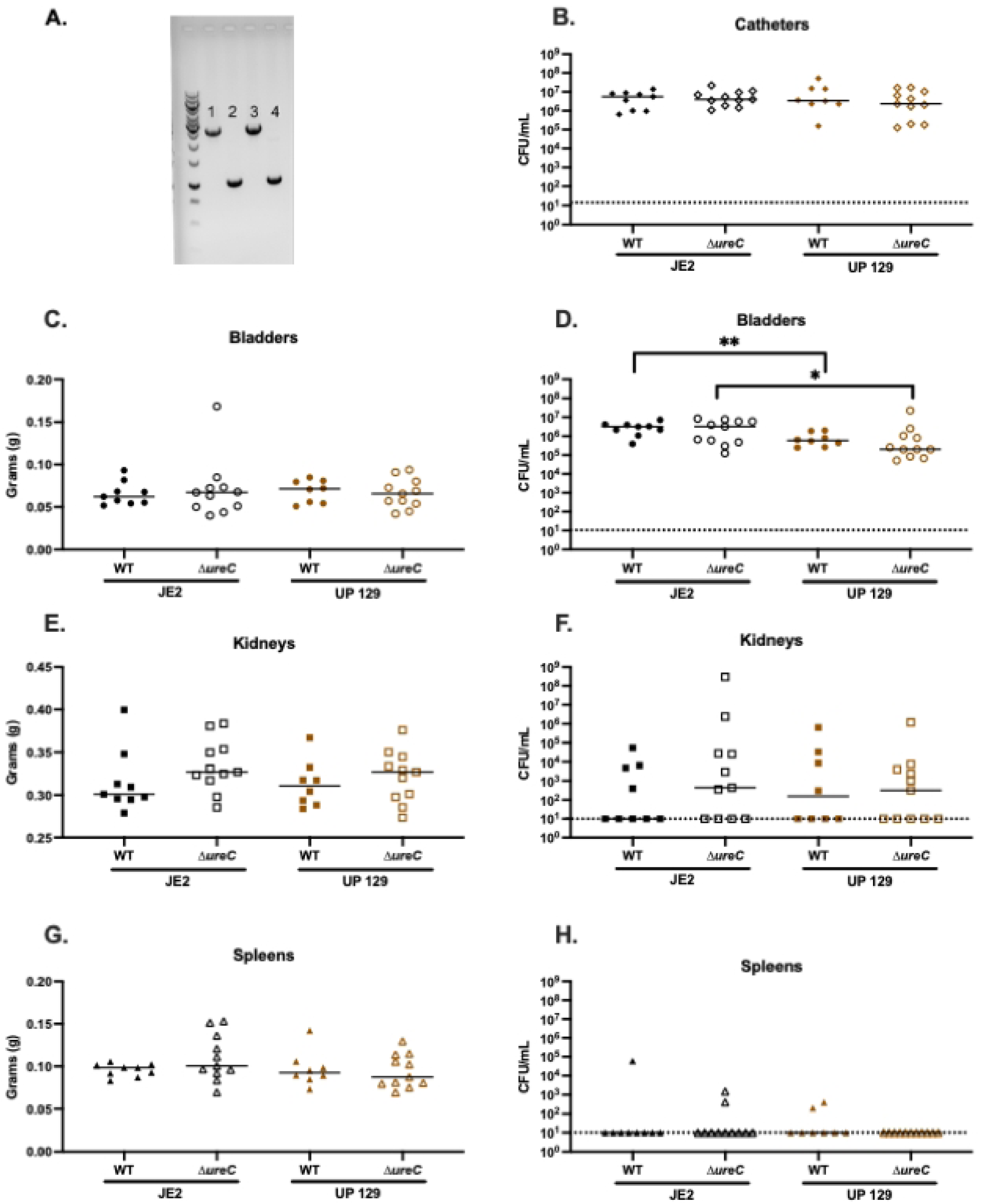
Isogenic *ureC* mutants assessed at 7dpi. **A)** PCR was performed using the ureC KO upstream F and ureC KO downstream R primers listed in Table 2. JE2 wild type (1) and UP 129 wild type (3) had large bands (∼1.7kb), indicating a fully intact *ureC* gene, while JE2 *ΔureC* (2) and UP 129 *ΔureC* (4) had smaller bands (∼0.5kb), indicating a deletion of *ureC.* **B-H)** CAUTI mouse model of *S. aureus* wild type and *ΔureC* strains. Mice were infected with either JE2, JE2 *ΔureC,* UP 129, or UP 129 *ΔureC* for 7 days. Colony forming units (CFUs) were obtained from the catheter (**B**), bladder (**D**), kidneys (**F**), and spleen (**H**). The weights of the bladders (**C**), kidneys (**E**), and spleens (**G**) were also measured at each time point. Each diamond, circle, triangle, and square represent one catheter, bladder, kidney, and spleen from an individual mouse, respectively. Closed black shapes indicate mice infected with JE2 and open black shapes indicate mice infected with JE2 *ΔureC.* Closed brown shapes indicate mice infected with UP 129 and open brown shapes indicate mice infected with UP 129 *ΔureC.* The Mann-Whitney U Test was used to determine statistical significance between groups, where *** is p<0.0005, ** is p<0.005, * is p<0.05, and p>0.05 is no significance. Solid lines indicate the median.

**S1 File:** Sequenced pUre plasmids.

**S2 File:** List of genomes used in this study.

